# A PDMS-based microfluidic platform enabling dual biochemical and electric-field stimulation for modeling sensory neuron-intervertebral disc crosstalk

**DOI:** 10.64898/2026.07.29.741573

**Authors:** Olanrewaju Akande, Sade W. Clayton, Liufang Jing, David Duong, Bryce Stottlemire, Ryan Potter, Mohammadjafar Hashemi, Addison Liefer, Nathaniel Huebsch, Lori Setton, Simon Y. Tang, Cory Berkland

## Abstract

Inflammation-driven increases in nociception are prominent in pain pathologies associated with intervertebral disc (IVD) degeneration yet are difficult to model in vitro. Since neurons are exposed to multi-modal stimuli in vivo, it is critical for these exposures to be conserved in an in vitro test system. We developed a polydimethylsiloxane (PDMS)-based microfluidic platform to interrogate peripheral sensory neurons (SNs) in the presence of conditioned media from nucleus pulposus cells from the degenerated IVD, to model a potential impact of IVD cells’ secretome on pain sensing. Our platform enables controlled perfusion of cell-derived biochemical cues alongside a defined homogeneous electric field (EF) and supports real-time optical analysis. Computational modeling, fluid perfusion experiments, and conductivity measurements confirmed stable fluid transport and tunable homogeneous EF generation within the device. As proof of concept for neuronal stimulation, neuroblastoma (N2a) cells loaded with a fluorescent Ca^2+^ indicator exhibited a 56% increase in Ca^2+^ transient activity when exposed to media from degenerated IVDs, concomitant with increased IVD-derived IL-1β production. Importantly, EF-stimulated Ca^2+^ transients increased in SNs derived from human induced pluripotent stem cells when exposed to conditioned media from primary human IVD cells, demonstrating the translation of this model system to human cells. Together, these results establish a versatile platform that enables controlled and simultaneous exposure to biochemical and electrical stimuli to quantify inflammation-driven peripheral neuronal hyperexcitability in tissue–neuron crosstalk.

## 1 Introduction

Lower back pain imposes an annual economic burden surpassing $100 billion USD in the U.S. with an average cost of $10,000 per patient^1,2^. These numbers encompass the direct and indirect costs that include initial diagnoses, treatments, and lifestyle changes. Tissue injury and inflammation promote downstream increases in nociception indicative of pain-like phenotypes. Increased nociception is also related to intervertebral disc (IVD) degeneration, spondylosis, stress fractures, and similar diseases where inflammation is central in pain-related pathophysiology.

However, isolating and studying neuronal hyperexcitability in vivo is difficult. Although in vivo animal models provide clinically relevant insights, they also present challenges such as ethical concerns, time consumption, and high costs that have prompted funding agencies to discourage extensive animal studies^3,4^. Traditional in vitro tissue culture models provide more control over confounding variables but often struggle to obtain real-time, biologically relevant pain-associated neuronal responses to inflammation like induced calcium (Ca^2+^) transient activity and proliferative changes, making it challenging to correlate experimental observations with clinical phenomena^5–9^. These limitations collectively highlight the need for alternative experimental platforms that can bridge the gap between simple in vitro cultures and complex in vivo systems.

Promising platforms for studying neuronal hyperexcitability that integrate sensory neuron (SN) subtypes are organ-on-chip (OoC) technologies^10,11^. OoC systems are microengineered to model specific in vivo phenomena experienced by tissues. These technologies provide the operational ease and high throughput inherent to traditional in vitro culture systems while supporting microenvironments that recapitulate crucial aspects of in vivo conditions like shear stresses from fluidic perfusion and system-wide electric stimulation. Additionally, several studies have highlighted the importance of emulating naive conditions and metabolite build-up to promote high nutrient concentrations^12–16^. Some neural OoCs specialize in maintaining chemical treatments within culture chambers while others focus on delivering electrical stimulation^17–19^. While these platforms represent improvements in this field, most fail to address key challenges associated with sustained neuronal cultures or present other drawbacks like a lack of generalizability and high complexity that do not justify transitioning from traditional in vitro cultures^17,19,20^.

Neurons also have biological and physical requirements that must be considered during device design. Their highly polarized morphology and need for high network organization necessitate large culture chambers to prevent loss of directional signaling and uncontrolled neurite growth. Next, their extreme sensitivity to shear stresses accompanying fluid exchange necessitates minimization of internal fluid-induced mechanical stress. Furthermore, neuronal cultures benefit from moderately dense environments promoting network interconnectivity and intracellular signaling, meaning large cell numbers must be accommodated within devices^17,21^.

Here, we designed a polydimethylsiloxane (PDMS)-based microfluidic platform through low-cost 3D-printed mold generation that facilitates tunable chemical and homogeneous electric field (EF) stimulation within a neuronal culture chamber. This field was achieved by lining opposing walls within the chamber with electrodes, resulting in a cylindrical homogeneous region spanning the chamber. The device’s large culture chambers also accommodated cell-seeded glass coverslips, transparent PDMS to enable optical analysis, and an internal geometry minimizing fluidic shear. Additionally, cells seeded on glass coverslips within the chamber could be electrically stimulated without removing them from the device, minimizing contamination risk. Computational simulations exploring fluid flow and homogeneous EF distribution validated the device’s viability. Initial EF stimulation studies with murine neuroblastoma (N2a) cells validated computational simulations, and we sought to capture differences in N2a excitability within biologically complex inflammatory environments. A similar human IVD cell co-culture model with induced pluripotent stem cells (iPSCs) was used to further evaluate hyperexcitability. In vivo hyperexcitability is difficult to quantify, so we address this by linking it to increased nociception through isolating IVD-neuron signaling within a controlled in vitro microenvironment. This analysis has been difficult in previous models because separating the native immune response, filtering desired signaling pathways, and other processes are required.

Prior work by our team identified Vascular Endothelial Growth Factor A (VEGFA) as a key indicator of IVD degeneration and that injured IVDs secrete inflammatory factors like IL-6, IL-8, and TNF-α which promote tissue degeneration and alter regional homogeneity^22–26^. Building on this, we hypothesized that SN-like N2a cells exposed to media secreted from injured IVD tissue would exhibit increased levels of EF-induced excitability. Then in our human model, we hypothesized that silencing VEGFA in upstream IVD cells before exposing their inflamed secretome to iPSC-differentiated SNs would decrease downstream hyperexcitability linked to pain-like responses. We uncovered links between IVD inflammatory signaling and neuronal excitation in both models, pointing to the significance of the IVD-SN signaling pathway. The various studies conducted within this device also highlight its potential as a versatile bioelectronic interface for neuronal inflammation and related pathological investigations.

## 2 Experimental

### 2.1 Device validation and characterization

#### 2.1.1 Design criteria

The microfluidic device was divided into 2 layers; the 5.4-mm bottom layer featured two 28-mm long, 0.4-mm wide electrode slots and two 16-mm square chambers connected by three 4-mm long, 2-mm wide channels, all with a depth of 3.4 mm in the bottom layer. The 2-mm thick top layer also had two 16-mm square openings that corresponded to the chamber positions in the bottom layer. Both layers also had a pair of 1.5-mm diameter hemisphere cylinders etched parallel to the device’s long side that supported tubing attachments. PDMS plugs matching the chamber geometries were created to seal the inner chambers when the device was in use (Figure 1).

**Figure 1:**
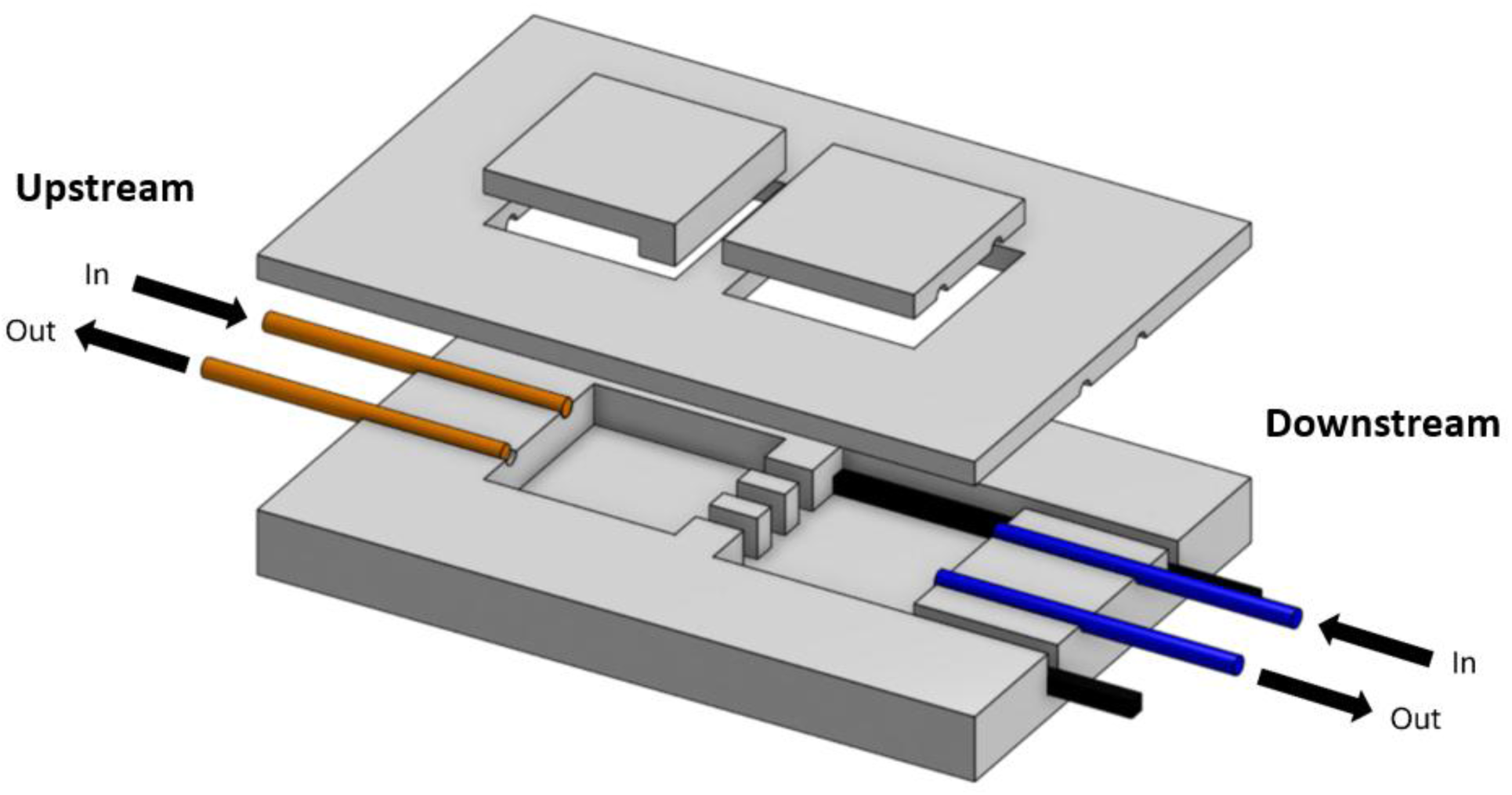
Microfluidic platform schematic detailing inner geometries. Two square culture chambers were linked by three channels. SS 316L electrodes line opposing walls of the downstream chamber and fluidic ports are located on both sides of the device. Plugs ensure the microenvironment is sealed.

#### 2.1.2 Computational simulations of fluid flow and electric fields

Fluid movement through the device’s bottom layer was modeled in COMSOL Multiphysics 6.3 because this was where stimuli were applied. During material selection, PDMS was assigned to the bulk bottom layer, stainless steel 430F to the two electrodes, and the inner regions were defined as water (Supplementary Tables 1-3). Solid mechanics, laminar fluid flow, and EF nodes were then applied to the appropriate nodes, and a “fine” mesh (small tetra- and hexahedra, pyramids, and prisms) was applied to the entire build before a stationary (time-independent) study was run (Figure 2). Incident flow velocity was controlled to assess affected pressure and shear throughout the device in single- and multi-inlet/outlet flow conditions (Figure 2, Supplementary Figure 1).

**Figure 2:**
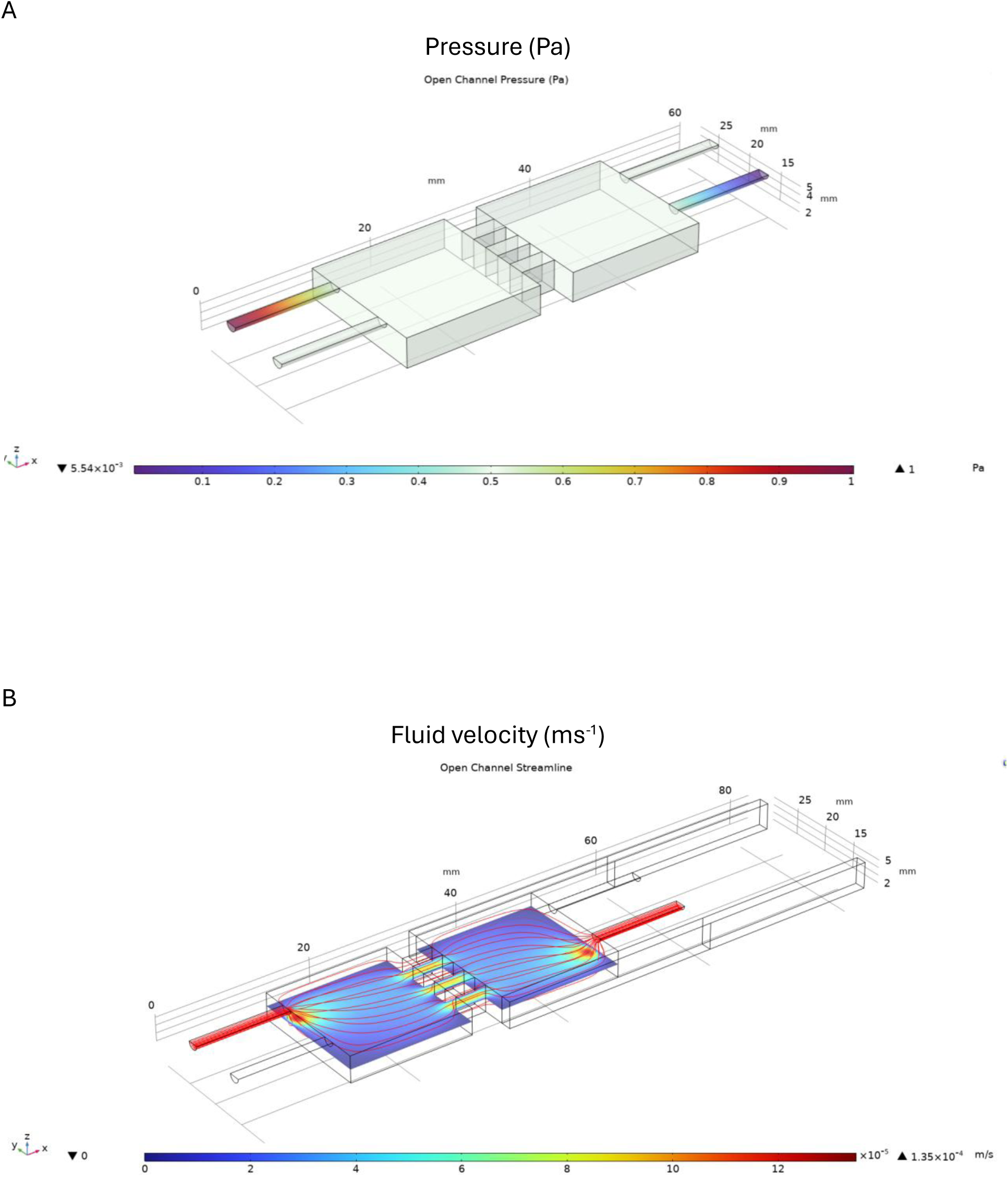
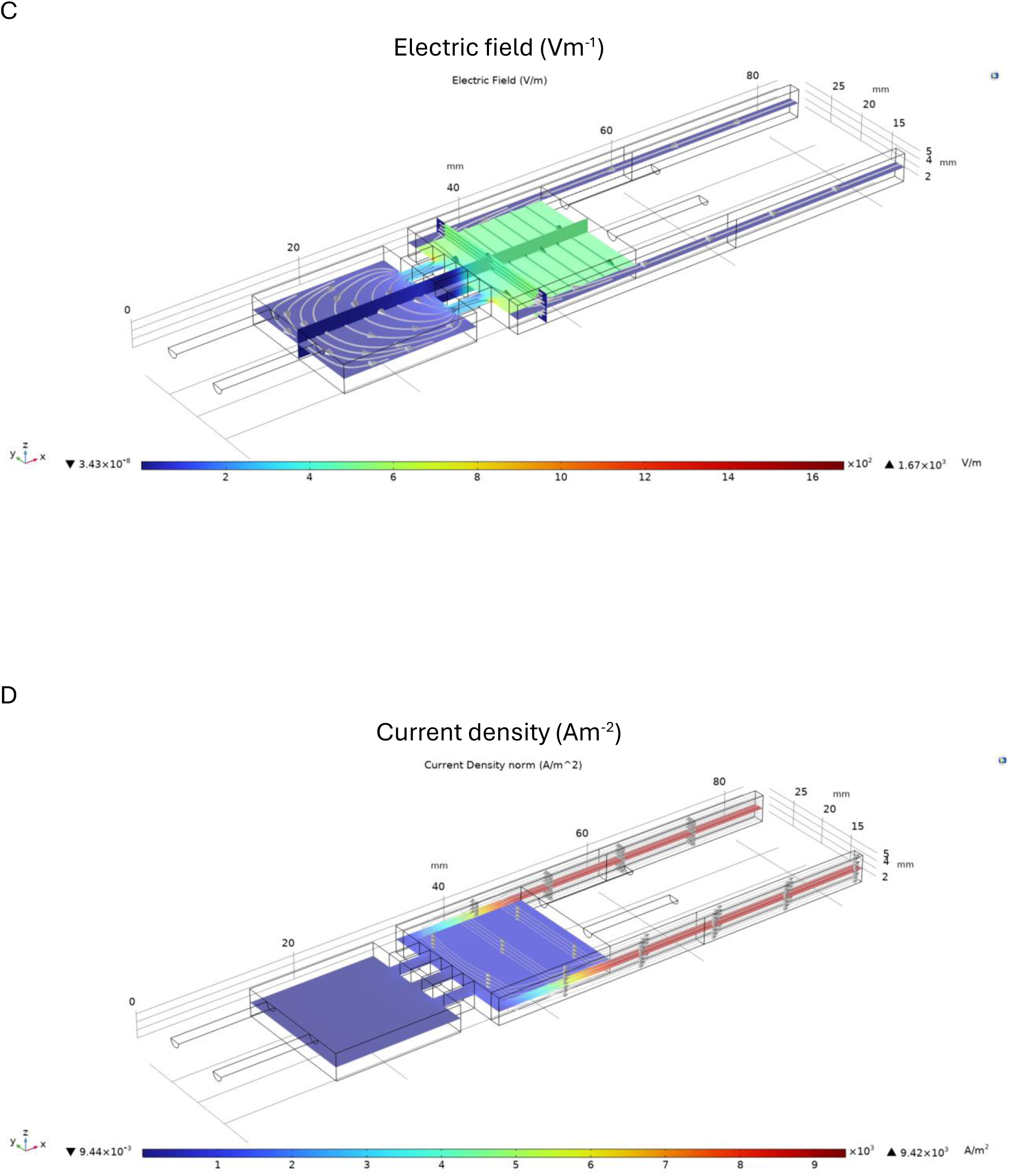
Microfluidic platform simulations assessed stimulation support. COMSOL Multiphysics 6.3 simulations covered pressure (**A)**, fluid perfusion **(B)**, electric field stimulation **(C)**, and current density field generation **(D)** in the downstream chamber.

#### 2.1.3 Manufacturing process

A double positive mold made from photopolymer resin (ELEGOO, Grey, 405 nm UV-curable, Guangdong, China) was fabricated through liquid crystal display printing with a 50 µm layer height to ensure precise geometry rendering. After a 10-minute wash in isopropyl alcohol, the double mold was cured under UV light (405 nm) for 10 minutes and sonicated in deionized water for half an hour. After drying, a 3% agar mixture was created and raised to 85°C to dissolve the agar in solution. The mixture was then poured into the resin mold to make an agar “negative” and left at room temperature for 30 minutes before being covered in Parafilm and transferred to 4°C storage for at least 2 hours.

Silicone elastomer curing agent (Dow Corning, Sylgard-184, Midland, MI, USA) and its corresponding base were gathered in a 1:10 ratio (w/w) before undergoing a 6-minute mixing and defoaming process in a Thinky mixer (Thinky USA, AR-100, Laguna Hills, CA, USA) at 2000 rpm. The mixture was then poured into the chilled agar negatives, de-gassed, and cured at 60 °C for 18 hours.

After curing, both layers were removed from their agar molds and sonicated in isopropyl alcohol for 30 minutes before being left to dry. The inner surfaces of both layers were corona-treated (Electro-Technic Products, BD-20, Chicago, IL, USA), pressed together, and cured at 60 °C for 2 hours. Stainless-steel 316 (SS 316L) wires (McMaster-Carr, 92705K145, Elmhurst, IL, USA) were then inserted into their corresponding slots in the device and the cured PDMS plugs were added to the top layer to complete the device.

Before being used, the device was disinfected by being submerged and sonicated in an ethanol bath for 1 hour. After sonication and subsequent evaporation of trace amounts of ethanol solution from within the device, the device was autoclaved at 121 °C and stored until its use.

#### 2.1.4 Validation studies

##### 2.1.4.1 Device trials

Two programmable syringe pumps (Harvard Apparatus, PHD 2000 Programmable, Holliston, MA, USA) were oriented to face each other with two microfluidic platforms between them (Supplementary Figure 2). Two 5 mL syringes held by syringe holders were placed onto the downstream pump with the syringe plungers completely compressed, followed by placement of 1.6 mm dia. polyethylene tubing line with each of the devices’ outlet ports. Two 5 mL syringes were then filled with 5% trypan blue and placed onto the upstream (left) syringe pump. Both pumps were then programmed to either infuse or refill media (upstream pump programmed to infuse, downstream pump programmed to refill) at a rate of 0.125 mL/ hour. Upon the program’s completion, devices were inspected for any leakage that occurred during the run. This run was repeated, and then 3 more iterations were done at rates of 0.625, 1.25, and 2.5 mL/ hour.

##### 2.1.4.1 EF stimulation

Alligator clips were attached to each of the electrodes within the device and plugged into a function generator (IONOPTIX, MyoPacer EP, Westwood, MA, USA). An additional electrode was placed into the device’s downstream chamber and attached to an oscilloscope. Then, a saline solution was perfused into the chamber, and a biphasic pulse (937.5 V/m, 200 ms/ on 1800 ms off) was sent through the device’s electrodes. Comparative analyses were performed between the field sent through the device’s electrodes and the field read through the additional electrode to confirm the field generated through the chamber.

### 2.2 Tissue and SN-like cell conditioned media experiments

#### 2.2.1 Cell source N2a cells

Mouse neuroblastoma cells (MilliporeSigma, N2a, SCC927, Burlington, MA, USA) were purchased and cryopreserved in liquid nitrogen prior to use in subsequent studies. Cells were allowed to attach to plasma treated surfaces of 12-well plates cultured in standard media (EMEM, 10% FBS, 1% penicillin-streptomycin) and incubated at 37 °C and 5% CO_2_. Cells were passaged every 3-4 days until used in the microfluidic platform for subsequent experiments.

#### 2.2.2 Organ culture of IVD and injury model

Eight virgin female 7 to 8-month-old mice (C57BL/6J) were euthanized within a CO_2_ chamber and sterilized with 70% ethanol solution prior to functional spinal unit (FSU) harvesting adhering to Washington University in St. Louis School of Medicine Institutional Animal Care and Use Committee guidelines. Three caudal (tail) FSUs were obtained from each mouse and wrapped in PBS-soaked gauze prior to tissue culture^8^. FSUs (CA1/2- to CA7/8) were carefully dissected and separately placed into a single well of a 24-well plate with 2 mL of culture media (DMEM + 20% fetal bovine serum (FBS) + 1% penicillin-streptomycin) and incubated at 37 °C with 5% CO_2_ and 100% humidity. Culture media was exchanged once daily for the next five days. On the sixth preconditioning day, wells were divided into four groups (G1-G4, n = 3 mice per group) that determined their injury treatment. The injury consisted of a bilateral needle puncture with a 30G needle through an FSU’s IVD. In G1, none of the FSUs were punctured; in G2, one FSU was punctured; in G3, two FSUs were punctured; and in G4, all the FSUs were punctured. After injuries, the FSUs were returned to culture in fresh media, with the media being collected and replenished after three days before ending the tissue culture after the sixth day post-injury. Collected media samples were stored at −80 °C to preserve media and cytokine integrity. An ELISA was used to assess IL-1β levels at the 3-day post-injury.

#### 2.2.3 Mouse neuroblastoma cell cultures

Neuroblastoma (N2a) cells were recovered from cryopreservation, cultured in standard media (EMEM, 10% FBS, 1% penicillin-streptomycin), and incubated (37 °C, 5% CO_2_, 100% humidity). After the second passage, cells were seeded on poly-d-lysine-coated 12 mm glass coverslips with a 50,000 cells/coverslip density. Standard media was given to the cells 24 hours before being replaced by differentiation media (EMEM, 1% FBS, 1% penicillin-streptomycin + 0.1% 10 μM retinoic acid) for the next 48 hours^27,28^. Coverslips were then randomly assigned to one of four treatment groups (n = 3) receiving a 1:1 mixture of differentiation media and injured murine FSU-generated media from groups G1- G4. Cells were cultured in this media mixture for 48 hours.

### 2.2.4 Mouse neuroblastoma cell stimulation and Ca^2+^ transient capture

The differentiated N2a cells were incubated in 3 µM of Fluo-4 AM (Thermo Fisher, reconstituted in DMSO), a fluorescent Ca^2+^ dye, in a Hank’s Balanced Salt Solution (HBSS) vehicle in darkness for 1 hour at room temperature. The HBSS media was then removed, and the cells were gently washed three times with fresh HBSS before being suspended in HBSS. The glass coverslips were then placed in the microfluidic device, submerged in HBSS, and positioned on a Nikon Eclipse Ts2r-FL microscope stage (Figure 3B). For chemical depolarization, a 40-second video was collected (50 ms exposure, frame rate of 20 frames per second) and a 30 mM KCl solution was added to the cells to induce Ca^2+^ transients. Electrically stimulated N2a cells underwent the same Fluo-4 AM and HBSS loading cycles before a function generator was attached to the device’s electrodes via alligator clips and positioned on the microscope stage. Each electrical stimulation trial ran for 40 seconds and was recorded with 50 ms exposure at a frame rate of 20 frames per second. Cells were stimulated with a biphasic source (30 V, 10 Hz frequency, 10 ms pulse width) for 5 seconds (frames 101-200) and allowed to rest for the remainder of the trial. Cell fluorescence was then analyzed in Fiji to track Ca^2+^ transients across cell membranes^29,30^. Fluorescence increases of at least 20% above baseline were defined as Ca^2+^ transient activation.

**Figure 3:**
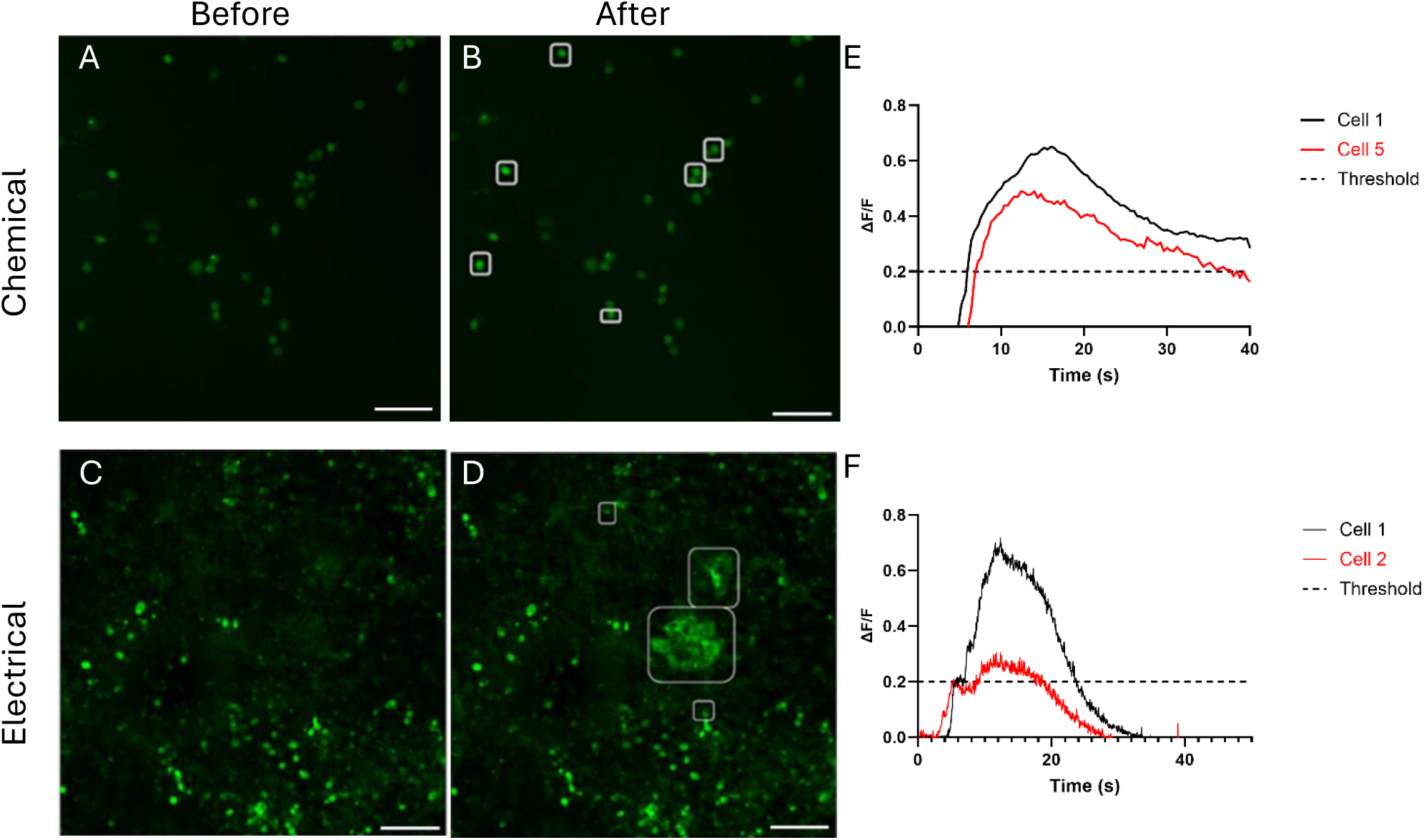
Chemically and electrically induced Ca^2+^ transients in mouse N2a cells. Images of the cell chamber before and after chemical (30 mM KCl) **(A-B)** and electrical (30 V biphasic, 5 ms pulse width, 5 Hz frequency) **(C-D)** stimulation enabled Ca^2+^ transient observation marked by the boxed cells. Fluorescent changes after chemical **(E)** and electrical **(F)** stimulation were quantified. Cells exhibiting increases of at least 20% in fluorescence intensity were surrounded by white boxes and classified as Ca^2+^ transient activity sites. Scale bar = 100 μm.

After this initial study was completed, another electrical stimulation regimen was performed with 100-second-long duration to test repeatable excitation. The EF stimulus and frame collection parameters were identical, but cells were now stimulated for two 10-second windows (frames 101-300 and 1501-1700) with a 60-second rest period between stimulation windows. Each coverslip was divided into thirds, and a field of view (FOV) (0.443 mm^2^) was captured within each region for a total of 3 fields collected per coverslip. A representative percentage was then calculated for each coverslip dictated by the number of responsive cells, C_r_, amongst viable cells, C_v_, in each FOV where the subscript 1, 2, or 3 corresponds to the FOV (equation 1).

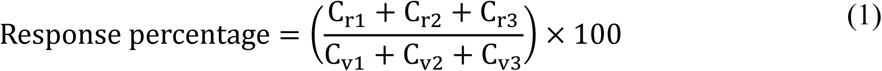

### 2.3 Human IVD cell and iPSC-derived sensory neuron culture condition

#### 2.3.1 Primary IVD cell conditioned media

Human donor IVD tissue (female, 50 and 70 y/o) was obtained freshly from the operating room as surgical waste. IVD tissue was immediately washed (HBSS, 5% FBS, 1% pen-strep), minced, and digested in Pronase (DMEM/F12, 5% FBS, 1% pen-strep, 0.2% Pronase) for 60 min at 37°C under gentle agitation. Samples were then removed from digestion, rinsed with HBSS (HBSS, 1% pen-strep), and digested in a collagenase solution (DMEM/F12, 5% FBS, 1% penicillin-streptomycin, 0.025% collagenase type II) overnight at 37°C. After digestions, IVD cells were separated from remaining tissue debris with a 70 µm cell strainer and then treated with RBC Lysis Buffer (10X) (Biolegend). Isolated cells were centrifuged, counted, resuspended in freezing media (90% DMEM/F12, 10% DMSO), and stored at −80 °C until needed. For these experiments, IVD cells were thawed and plated in a 75-cm^2^ culture flask with culture medium (DMEM/F12, 10% FBS, 1% penicillin-streptomycin) to expand to 90% confluency in a humidified incubator (37 °C, 5% CO_2_, 20% O_2_). Cells were detached with 0.05% trypsin (trypsin/ethylene diaminetetraacetic acid) and re-plated (3 x 10^5^ cells/mL) in 175-cm^2^ culture flasks with 2 mL of IVD medium and allowed to reach 80% confluency.

IVD cells were then plated into four wells of a 6-well plate (300,000 cells/well) and randomly assigned an siRNA transfection and IL-1β stimulation treatment (n = 3). Each well either served as a no stimulation control, an IL-1β stimulation only well (1 ng/mL), a scrambled siRNA-GFP (sc-36869, Santa Cruz Biotechnology, Dallas, TX, USA) + IL-1β stimulation well, or VEGFA-siRNA (sc-29620, Santa Cruz Biotechnology, Dallas, TX, USA) + IL-1β stimulation well. First, the IVD cell culture medium was replaced with OptiMEM™ and cells in the designated wells were transfected with 30 pM of scrambled siRNA or VEGFA siRNA via utilization of Lipofectamine RNAiMax™ (cat # 13778100) reagent for 24 hours. Transfection media was removed and replaced with IVD cell culture media and images were taken of the GFP-tagged scrambled siRNA to observe the cell transfection percentage. Cells were allowed to rest for 24 hours before stimulation with IL-1β in the designated wells. After 24 hours, the IL-1β stimulation media was exchanged for fresh IVD cell culture media and incubated for 48 hours. Then, media from each well was collected to assay human VEGFA via ELISA (Mouse VEGF, DY293B, R&D Systems, Minneapolis, MN, USA) to determine the efficiency of VEGFA knock-down. The cells were lysed in 300 mL of TRIzol™ solution and placed in −80 °C prior to their application with the mature sensory neurons as described here. Materials and reagents were purchased from Thermo Fisher Scientific (Waltham, MA, USA) unless otherwise stated.

#### 2.3.2 Differentiation of iPSCs into sensory neurons

Human iPSCs (Wild-type Conklin Lab; Coriell repository #GM25256) carrying a genetically encoded Ca^2+^ indicator (GCaMP6f) were seeded in a 6-well plate (80,000 cells/cm^2^) and cultured in neuronal crest differentiation medium (mTeSRTM1) before differentiation into the neural crest cell phenotype^9,30^. After six days of culture with daily full medium changes, cell differentiation into neural crest cells was confirmed via expression of the CD271 marker (Supplementary Figure 3). These neural crest cells were then plated on glass coverslips (12-mm, pre-coated from EMS GG-12-1.5-PRE, 60,000 cells/well) and overlaid with SN differentiation media to promote the SN phenotype. This differentiation media was replenished daily from days 7 to 13 before SN differentiation was confirmed via optical analysis. The differentiation media was then exchanged for SN maturation medium from days 14 to 19 with media replenishment every 2 days. Neuronal maturation was confirmed on day 20 by abundant expressions of TUJ1, peripherin, BRN3A, and ISL1 via immunostaining (Supplementary Figure 4). SN maturation media exchanges continued every 2 days for sustained cultures.

#### 2.3.3 Human iPSC derived sensory neuron stimulation

Neuronal maturation media was removed from mature sensory neurons and replaced with depleted maturation media (identical mixture without BDNF (brain-derived neurotrophic factor) and GDNF (glial cell line-derived neurotrophic factor)) for a 24-hour culture. Depleted media was then removed and replaced with a fresh 1:1 mixture of depleted maturation media and vehicle (n = 4), IL-1β-stimulated (n = 4), or IL-1β -stimulated with scrambled (n = 4) or VEGFA (n = 4) siRNA pre-treated donor IVD-generated media. Neurons were cultured in this media for two days prior to it being removed and replaced with HBSS and subsequent EF stimulation (Supplementary Figure 4). The experimental protocol for EF stimulation was like that described for the N2a exposure as described above, but with different field parameters (30 V biphasic source, 5 Hz frequency, 5 ms pulse width).

## 3 Results

### 3.1 Perfusion and EF testing validated COMSOL simulation results

Computational simulation of the device’s geometry and environment were simulated prior to fabrication to verify fields for homogeneity of fluid flow and EF across the chamber. The simulation focused on fields in the downstream chamber as that is where functional readouts were to be collected. Simulation of single inlet/outlet fluid flow through the device with simultaneous EF generation gave rise to a fluid velocity field with magnitude (sub-6 × 10^-4^ ms^-1^) and negligible pressure drop. Generated EF and current density were also bounded with less than 5% deviation from mean values over the 16-mm-long region between electrodes in the downstream chamber (Figure 2). Fluid current paths were also predicted to occur within the downstream chamber and could be altered when the number of channels between upstream and downstream chambers was changed from one to four channels. Rerunning the simulation after “blocking” the channels between the chambers gave rise to nearly equivalent magnitudes and uniformity of pressure, fluid flow, and current density across the downstream chamber (Supplementary Figure 1). The primary difference between the two conditions was that the EF was denser in the closed channel condition even though the field’s magnitude was like the open channel condition (Figure 2C, Supplementary Figure 1C). This suggested the chambers’ geometries supported the design’s feasibility. These simulations supported the device’s design goal to ensure most of the chamber would experience a homogeneous EF and supported moving to fabrication of the physical device platforms.

Fluid perfusion testing with the fabricated device supported results of the pressure and fluid velocity simulations as fluid travelled along predicted routes and was ejected from the device as designed (Supplementary Figure 2). Similarly, EF testing was performed with biphasic pulses sent through the device’s electrodes and measurements of field strength were recorded. This showed there was a negligible difference (< 0.001%) between the EF sent into the device and the field measured within the chamber. Successful validation of stimuli facilitation encouraged progression into neuronal cultures within the platform.

#### 3.1.1 N2a cells were successfully depolarized through chemical and EF stimulation

Two methods of N2a stimulation were tested to see if they could prompt depolarization. First, chemical depolarization was tested through perfusion of KCl (30 mmol/L) in the upstream chamber, then, electrical stimulation was tested with an applied EF (937.5 V/m). Using the Fluo-4 AM fluorescent Ca^2+^ dye, plated N2a cells in the downstream chamber exhibited Ca^2+^ transient activity after KCl was perfused into the upstream chamber followed by a slow return to baseline values (Figure 3A,B). Applying an EF to the downstream chamber also drove Ca^2+^ transient activity associated with repolarizations, or a return to baseline values that were nearly 33% faster than recovery from KCl stimulation (Figure 3E,F). Effective N2a stimulation under baseline conditions gave proof-of-concept for the applicability of the device and was followed by stimulation under inflammatory conditions as described next.

### 3.2 Exposure to injured mouse IVD media increased responses in N2a cells

The concentration of IL-1β secreted from injured IVD motion segments was measured after periods of in vitro culture. This “conditioned media” was collected and perfused into the upstream chamber for tests of an effect on plated N2a cells exhibiting induced Ca^2+^ transients in the downstream chamber. When all three IVDs were injured in a group, there was a 1.14-fold increase in levels of IL-1β secreted over the control group (p < 0.05) (Figure 4A). Similar trends were also observed for IL-1β secreted values in groups in which only one or two IVDs were injured (1.04- and 1.06-fold). Perfusion of conditioned media from the three-injured IVD group was associated with a 14% increase in the percentage of cells in the downstream chamber exhibiting Ca^2+^ transients over those from the control group (p < 0.05) (Figure 4B).

**Figure 4:**
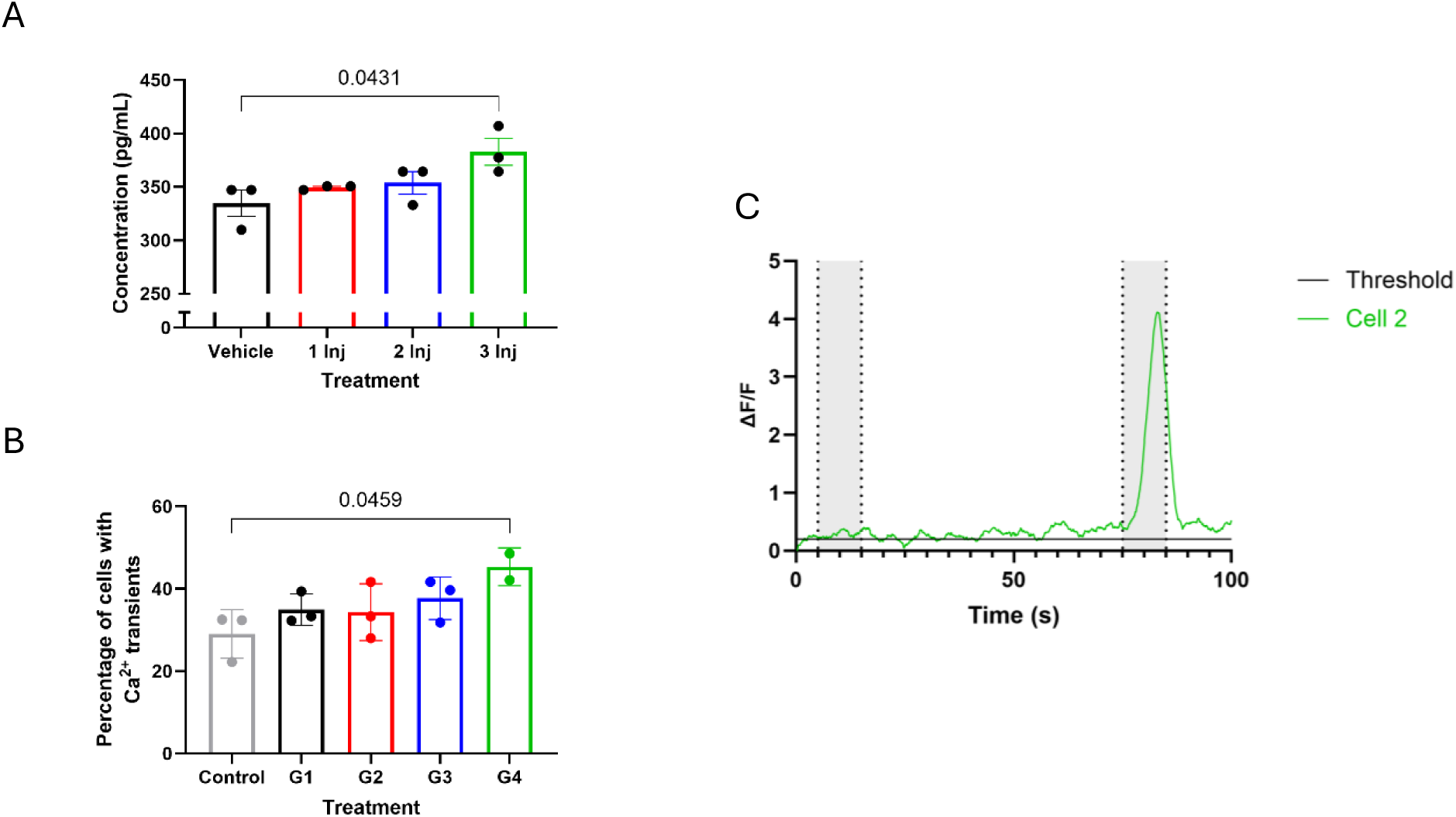
Increasing the number of injured murine IVDs led to increases in IL-1β concentration and average number of induced Ca^2+^ transients observed. ELISA readout of IL-1β concentration (pg/mL) produced by IVDs 3 days post-injury **(A)**. Percentage of cells responding to electrical stimulation after IVD media pre-treatment **(B)**. N2a neuronal groups cultured in media generated from zero (G1), one (G2), two (G3), or three (G4) injured FSUs in the same well. A representative cell **(C)** expressing fluorescent response to EF stimulation. Cells were stimulated twice for 10 seconds (grey regions) and rested for 60 seconds between the stimulation periods (white). Ca^2+^ transient activity modelling cell excitation was induced during stimulation periods. Experiment timeline can be found in Supplementary Figure 7.

### 3.3 Silencing VEGFA in human primary IVD cells diminished their ability to provoke Ca^2+^ transients in iPSC-derived SNs

Next, VEGFA’s role in content of secreted media from inflamed IVD cells was assessed, along with exposure of human iPSC-derived SNs to this conditioned media. In this experiment, SNs were exposed to media generated by IVD cells incubated with IL-1β. There was a 14% increase in the number of observed Ca^2+^ transients after EF stimulation compared to SNs exposed to conditioned media from IVD cells without IL-1β exposure. However, pre-treating IVD cells with VEGFA-silencing siRNA before IL-1β incubation modified the conditioned media produced, such that perfusion with this media decreased the number of observed Ca^2+^ transients in SNs by 75% compared to IL-1β incubation without siRNA (p < 0.05) (Figure 5A-B, Supplementary Figure 5). When normalized to the control group, directly inhibiting VEGFA production in IVD cells led to modified conditioned media conditions, associated with a 72% reduction in Ca^2+^ transients for SNs; in comparison, conditioned media from human IVDs treated with scrambled siRNA led to a 23% reduction in Ca^2+^ transients in plated SNs (p < 0.05) (Figure 5C).

**Figure 5:**
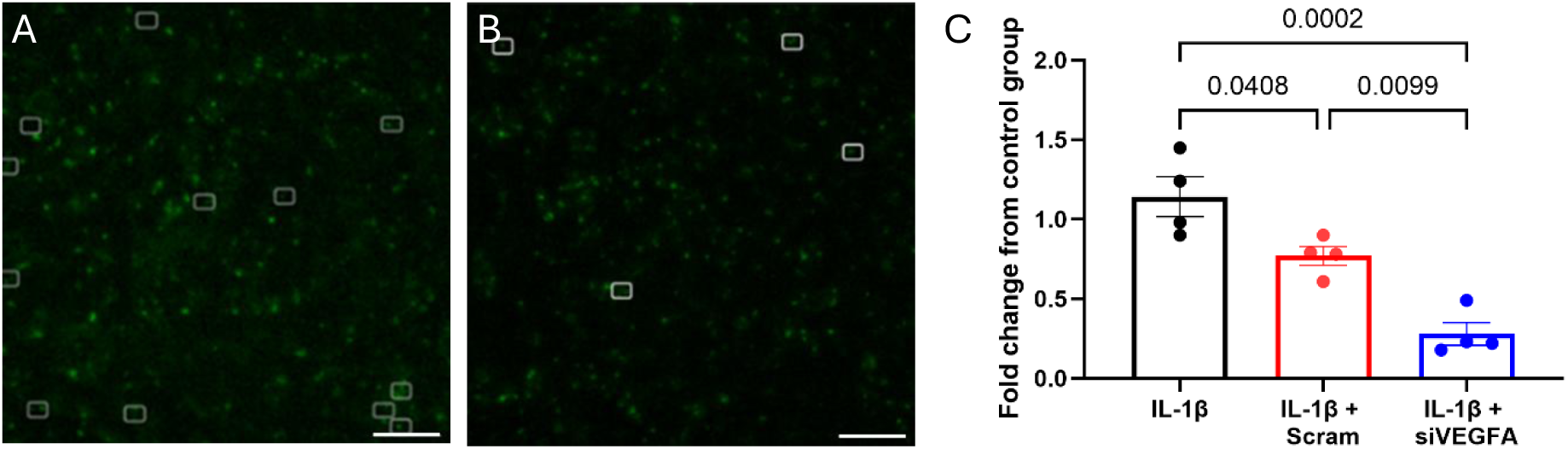
Human iPSC-derived sensory neurons exposed to media generated from VEGFA-silenced IL-1β -stimulated human IVD cells exhibited lower levels of induced Ca^2+^ transients. FOVs with similar cell densities captured neurons cultured in IL-1β-stimulated IVD media without **(A)** and with **(B)** the VEGFA-silenced siRNA pre-treatment prior to EF stimulation. Cells exhibiting induced Ca^2+^ transients are highlighted in white boxes (FOVs captured at 10X magnification). Fold change of each treatment normalized to the control group. Each data point represents a coverslip in which multiple FOVs were obtained **(C)**. Scale bar = 100 μm. Differentiation timeline can be found in Supplementary Figure 5A.

## 4 Discussion

Inflammation-driven hyperexcitability is key in many musculoskeletal diseases including IVD pathology, and in vitro model systems are useful for revealing the driving pathophysiologic mechanisms. This study focused on designing a microfluidic platform capable of providing dual biochemical and EF stimulation to monolayer cultures of excitable cells, such as neurons, with optical readout of Ca^2+^ activity. Computational simulations and physical model validation confirmed the device’s stimulus-delivering capabilities. Using it, the inflammatory IVD secretome prompted increases in neuronal excitability. Silencing VEGFA in human IVD cells exposed to the inflammatory stimulus, IL-1β, decreased downstream neuronal excitability. These findings demonstrate one utility for the platform that is supportive of mechanistic investigations into co-stimulation of cells affected by inflammatory stimuli in IVD disease.

Designing a platform for neuronal culture necessitated addressing a few criteria. While immortalized cell lines, like the N2a studied here, commonly divide, primary neurons and those derived from iPSCs are non-dividing, require weeks of culture for neurite outgrowth and expansion (maturation), and are susceptible to microenvironmental fluctuations^17,21^. The protocol for producing these cells also requires long cell timeframes, and the process itself is sensitive to the cellular environment. These factors would make it challenging to differentiate the iPSC-SN in situ within the device. The device was designed to accommodate these challenges by engineering a large downstream culture chamber with intentional internal channel geometries to support placement of attached SNs to prescribed substrate conditions while allowing for some additional neuronal network expansion. COMSOL simulations predicted laminar flow through the device and with velocities unlikely to promote detachment, as well as homogeneous EF generation in the downstream chamber. Exploratory studies with physical models confirmed these findings and neuron-seeded glass coverslips also supported real-time optical analysis of induced Ca^2+^ transients.

Many platforms support EF stimulation, fluid perfusion, and biochemical treatment, but few have demonstrated use with neuronal cultures^13,15,17,21^. Pavesi and co-workers designed a PDMS-based OoC device that delivered simultaneous electromechanical and biochemical stimulation to plated neuronal cells^15^. Although they were able to incorporate three types of stimuli into their device with homogeneous EF generation, their design relied on direct culture of cells onto PDMS. PDMS is not ideal for direct neuronal cell attachment as it absorbs culture media proteins, can leach toxic non-crosslinked oligomers, can undergo hydrophobic recovery over time, and absorbs hydrophobic small molecule drugs, making therapeutic concentration identification difficult^19,20,22^. We addressed these limitations by pretreating PDMS and using glass coverslips as the neuronal culture substrate. Commercially available perfusion chambers (e.g., Warner Instruments’ RC-49MFSH) have been fabricated to enable dual biochemical and electric stimulation but we found them unsuitable to support the long-term periods of culture needed to allow for neuronal cell maturation and branching prior to stimulation^31^. Our use of a closed system reduced the contamination risk and supported long-term culture, enabling our study of the same neuronal populations over different times and treatments. This negates the need for intragroup pooling across multiple neuronal systems that are sensitive to slight environmental changes, setting it apart from existing platforms (Table 1) ^31–34^.

**Table 1:**
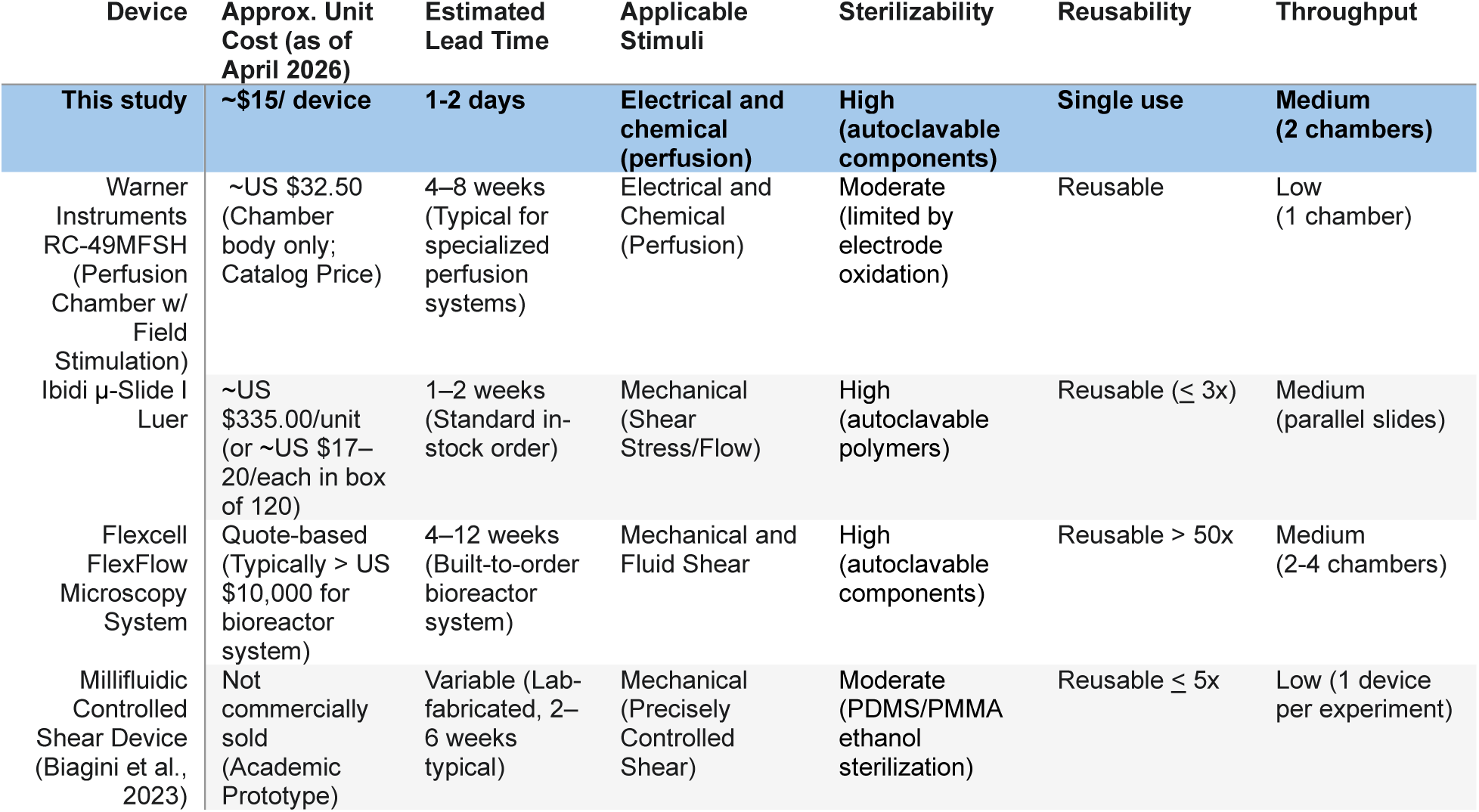
Comparing various microfluidic platforms’ unit cost, lead time, and key use cases to this study’s platform. This study’s device features a lower unit cost and faster lead time while facilitating multiple stimuli like other devices. PMMA- Poly(methyl methacrylate).

The device’s applicability for the study of neuronal crosstalk with chemokines released from murine IVDs proved to be of interest. Previous IVD studies have reported increases in inflammatory cytokine production after injury including elevated production of VEGFA isoforms and their receptors^7,22–26^. Increases in IL-1β secretion after IVD injury were measured here and are consistent with prior findings, although the magnitude of the increase over control was modest (p < 0.05). Nevertheless, we found exposing human iPSC-derived neuronal cells to injured IVD media prior to EF stimulation significantly increased Ca^2+^ transient activity (p < 0.05) suggesting that cytokine release following IVD injury was sufficient to drive neuronal excitability. This supports links between downstream increases in neuronal excitability after exposure to the injured and inflamed IVD secretome. It also revealed the successful isolation of the IVD-neuron pathway from confounding variables like the host’s immune response and injury application in vivo^35–37^.

Prior work had identified VEGF as a pivotal chemokine mediating vascular and neuronal ingrowth in IVD degeneration^8^. Potter and co-workers also reported a link between VEGFA production and pain related performance and sensitivity in mouse models, with findings of improved performance when VEGFA was ablated before injury in an in vivo murine model^8^. Our study of human IVD cell-conditioned media’s impact on human iPSC-derived SNs adds further to an understanding of these effects, as neuronal populations exposed to media from VEGFA-silenced IVD cells exhibited fewer Ca^2+^ transients than non-inflamed controls when interrogated by an EF (p < 0.05). Future studies can be performed to probe the composition of the IVD secretome and to better understand siRNA knockdown and IL-1β effects on VEGFA receptor expression to further investigate this signaling pathway in vitro using this microfluidic platform.

While the device design used here provided plating of neuronal cell populations and exposure to collected IVD secretome, the absence of direct contact between IVD tissue and neuronal cells limited the interpretation of co-culture conditions here. A true co-culture experiment would enable real-time analysis into the temporal nature of the inflamed IVD secretome as has been shown for studies of dynamic IVD culture systems that collect or process secreted IVD chemokines over time^38,39^. It could be instructive, for example, to understand how the injured IVD affects neuronal cell populations acutely and over long periods of time where cell adaptation has been permitted to occur. Another limitation during this process was that computational simulations assumed each of the tissue’s material properties were homogeneous. This assumption simplified the simulations and calculations, but other components like cell membranes, cytoplasmic properties, and extracellular matrix structure were not considered. Also, as a two-component device, the design will retain additional uncertainty concerning chamber isolation when compared to single-component devices. Furthermore, in vitro neuronal cultures are extremely difficult to maintain, and primary cells introduce another layer of complexity as the time from harvest to culture becomes a significant hurdle to overcome^40^. Although the large chamber dimensions support network formation, they do so at the expense of a lack in polarity enforcement that a micropatterned seeding substrate would provide. Despite these limitations, our device demonstrated its potential to support mechanistic studies into neuronal hyperexcitability that are difficult to achieve in existing models (Table 1).

## 5 Conclusion

This work describes the design of a new microfluidic device facilitating biochemical and homogeneous EF stimulation for multiple cell systems. Here the device was used to study IVD-SN signaling crosstalk pathway in an isolated and controlled manner, to investigate inflammation-prompted neuronal hyperexcitability. We show that media collected from injured mouse IVDs increases the number of EF-induced Ca^2+^ transients in human SN-like cells and that reducing VEGFA in human IVD cells prior to inflammatory challenge reduces downstream excitability. These results identify VEGFA as a potential contributor in IVD-SN crosstalk. Broadly, this platform can serve as a preclinical screening tool for anti-inflammatory therapeutic candidates targeting neuropathic sensitization in chronic degenerative and inflammatory disorders.

## 6 Author Contributions

**O.A.:** Conceptualization, Data curation, Formal analysis, Investigation, Methodology, Software, Validation, Visualization, Writing – original draft

**S.W.C.:** Investigation, Methodology, Validation, Writing – review & editing

**L.J.:** Investigation, Methodology, Validation, Writing – review & editing

**D.D.:** Data curation, Formal analysis, Investigation, Validation

**B.S.:** Conceptualization, Funding acquisition, Methodology

**R.P.:** Investigation, Methodology, Validation

**M.H.:** Investigation

**A.L.:** Investigation, Methodology, Validation

**N.H.:** Methodology, Resources

**L.A.S.:** Funding acquisition, Methodology, Resources, Writing – review & editing

**S.Y.T.:** Conceptualization, Funding acquisition, Methodology, Project administration, Resources, Supervision, Writing – review & editing

**C.B.:** Funding acquisition, Project administration, Resources, Supervision, Writing – review & editing,

## 8 Data availability

Raw and processed data supporting the findings of this study including computational simulations and quantified calcium transients are openly available in the Zenodo repository at https://doi.org/10.5281/zenodo.21477927.

## 9 Acknowledgements

## Acknowledgements

The authors would like to thank the Huebsch laboratory at Washington University in St. Louis for providing access to a Nikon Ts2r-FL microscope for Ca^2+^ transient analysis and the gift of the GCaMP6f-hPSCs.

## 13 Supplementary Information

### 13.1 Theory/Calculation

Theoretical formulations describing chemical transport and electrical stimulation within COMSOL Multiphysics were developed to guide device design and ensure controlled simulation conditions. These models informed the selection of stimulation parameters and provided a framework for interpreting experimental observations.

#### 13.1.1 Electrical stimulation

When the device was solely used as an electrical stimulation chamber, a potential formulation equation describing the system was sufficient. Initial assumptions that were made to direct equation derivation:

1. Stimulation occurs in stationary fluid
2. Stimulation media is water (with ions)
3. Quasistatic EF (time-independent)
4. Uniform material conductivity in the chamber and among the electrodes
5. Steady-state conduction with no net charge accumulation (charge conservation)

Boundary conditions were then added to further simplify successive calculations:

1. One electrode held at a fixed potential (+V)
2. One electrode held at ground (0 V)
3. Insulating PDMS walls with no current flux across the boundary

We then shifted to working with initial equations for the EF vector (E) and current density vector (J) within the defined constraints:

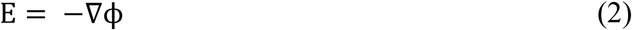

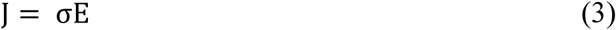

where ∇ϕ is the gradient of the potential and σ is the electrical conductivity of the medium. Substituting equation 2 into equation 3 results in another form to represent the current density vector, J.

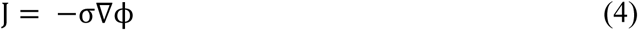

Applying the charge conservation assumption to our current density vector, equation 5 describes the lack of divergence of the current density vector. Rewriting equation 5 and simplifying while also accounting for the constant material conductivity results in equation 7 which is the original potential formulation equation simplified to Laplace’s equation.

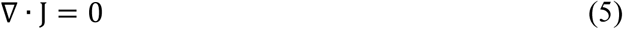

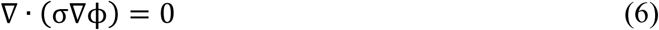

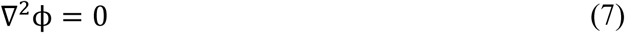

#### 13.1.2 Chemical stimulation

When the device was used for chemical stimulation without an electric stimulus, similar assumptions were made about the system:

1. Stationary fluid during chemical stimulation
2. Base media is water
3. KCl is treated as a single species
4. No flux at PDMS walls
5. Uniform initial concentration

Using these assumptions, the following boundary conditions at the pipette contact region were made:

1. C (t = 0) = C_0_
2. KCl characteristic diffusion

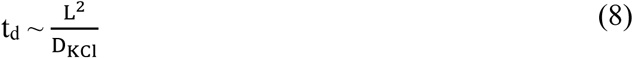

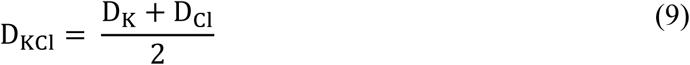

where t_d_, the characteristic diffusion time for KCl is approximately equivalent to the distance KCl diffuses squared divided by the diffusion coefficient of KCl in water. Equation 11 then links this to cellular depolarization as the increase of extracellular potassium ions, [K^+^]_e_, increases the potassium Nernst potential, E_K_, after KCl is added into the system.

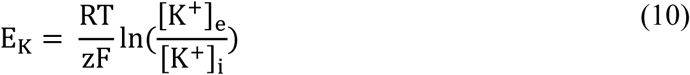

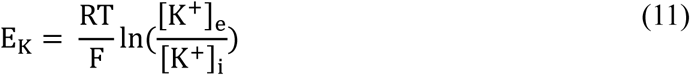

This equation shows how the increasing [K^+^]_e_ expands the difference between it and the intracellular potassium concentration, [K^+^]_i_, while also accounting for the effects of the universal gas constant, R, the system’s absolute temperature, T, Faraday’s constant, F, and the ion’s valency, z (for K^+^, z = 1).

### ***13.1.3*** Electrical stimulation with chemical stimulation

Combining both these scenarios together to define the dual stimulation model, assumptions must first be made that the species is migrating into the field:

1. Nernst-Planck (no flow)
2. Bulk electroneutrality

Assuming a Nernst-Planck scenario permits us to derive equations 12 and 13:

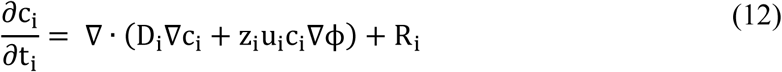

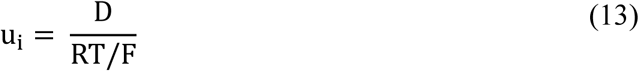

where c_i_, the species concentration’s change over time, t_i_, is a function of its diffusion coefficient, valency, concentration gradient, and electrophoretic mobility, u_i_. Our assumption for bulk electroneutrality with small concentration gradients ties back to equation 6 and enables the derivation of equations 14 and 15 detailing the species concentration’s effect on media conductivity using the Nernst-Einstein relation.

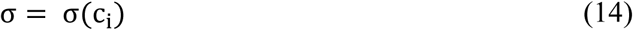

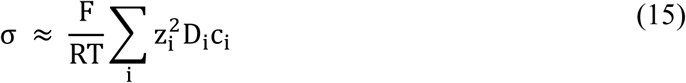

## 13.2 Supplementary Figures and Tables

**Supplementary Figure 1:**
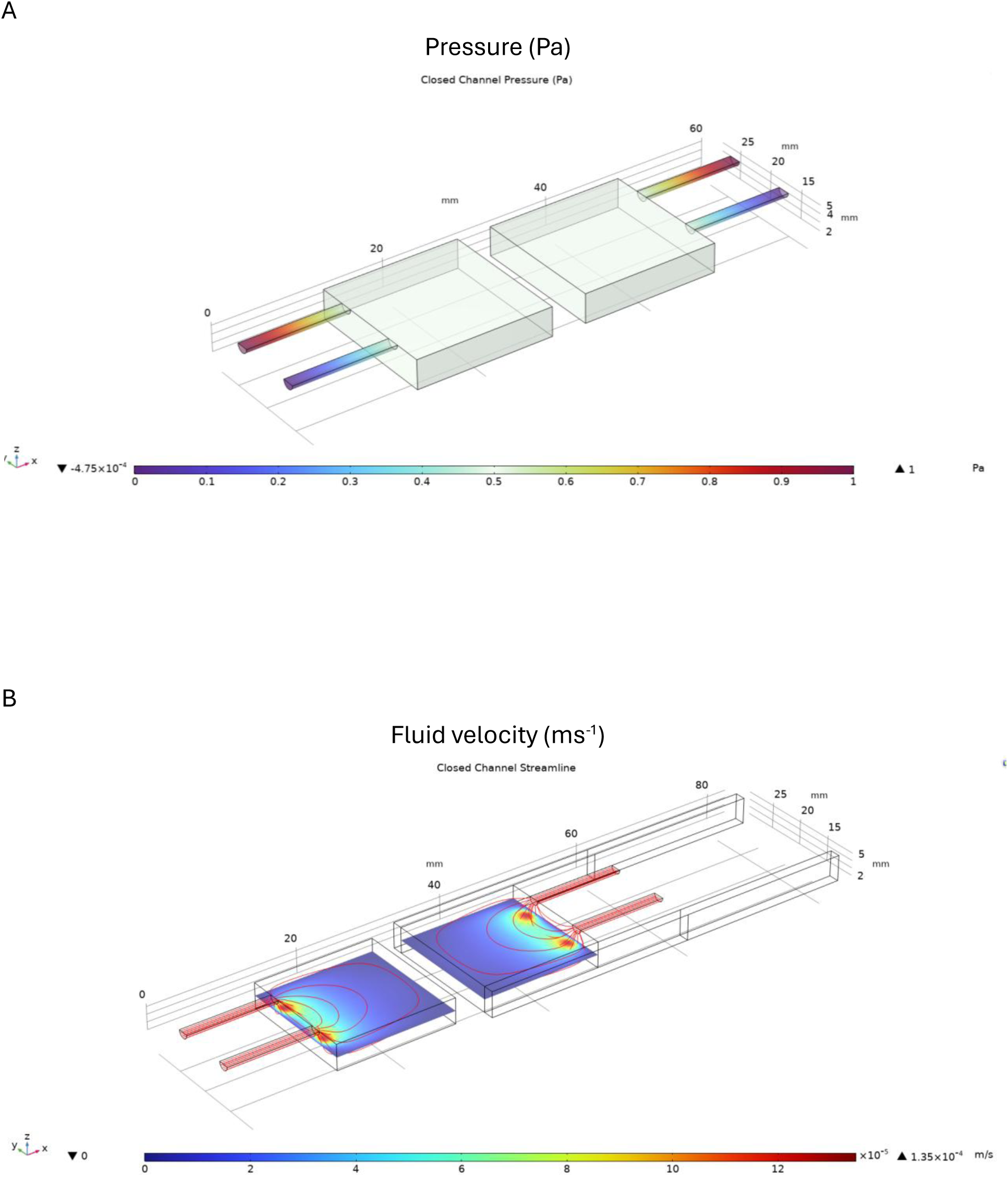
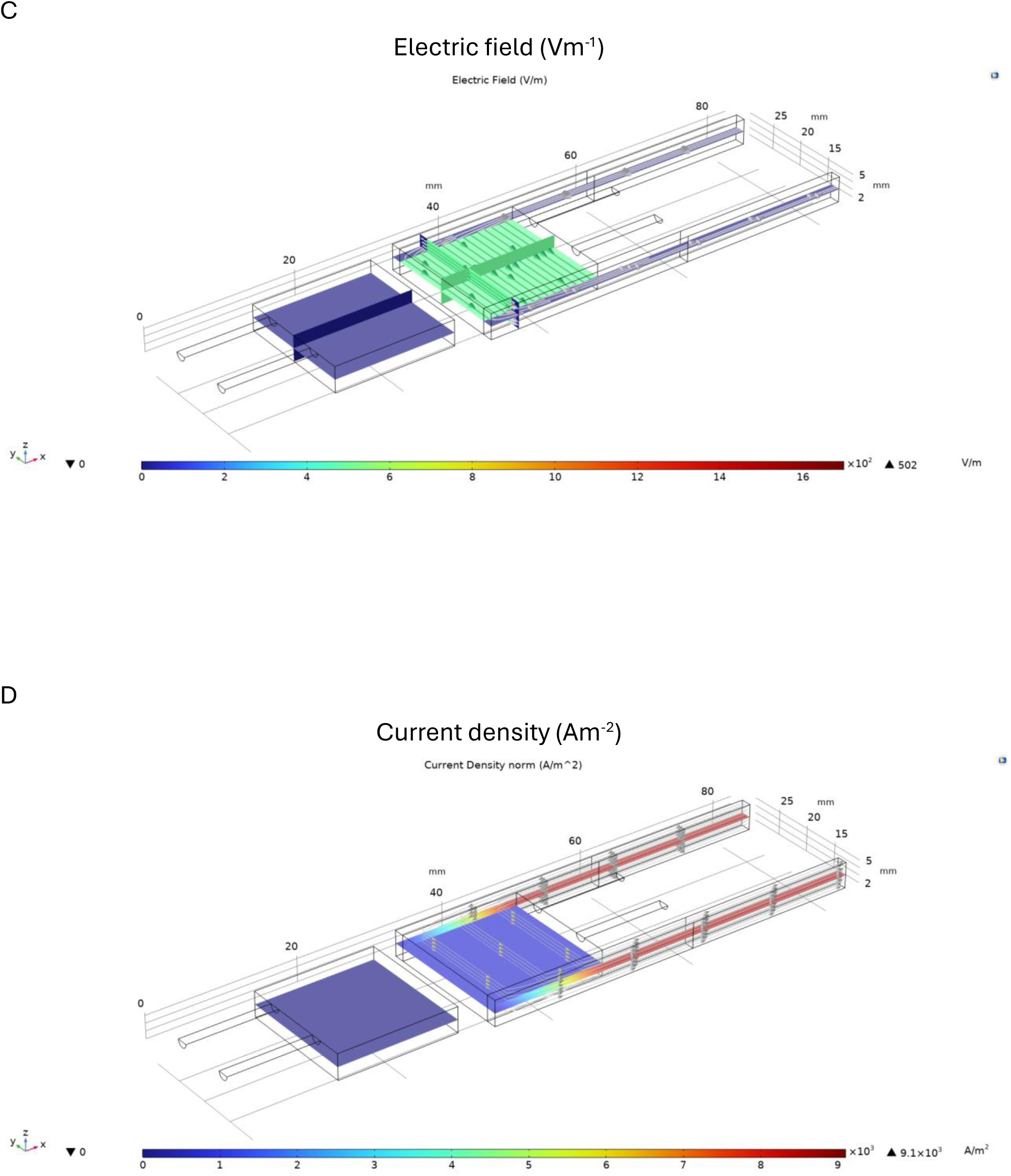
Microfluidic platform simulations with closed channels between chambers. COMSOL Multiphysics 6.3 simulations were performed in the closed-channel apparatus to assess differences in pressure (A), fluid velocity (B), electric field (C), and current density (D).

**Supplementary Figure 2:**
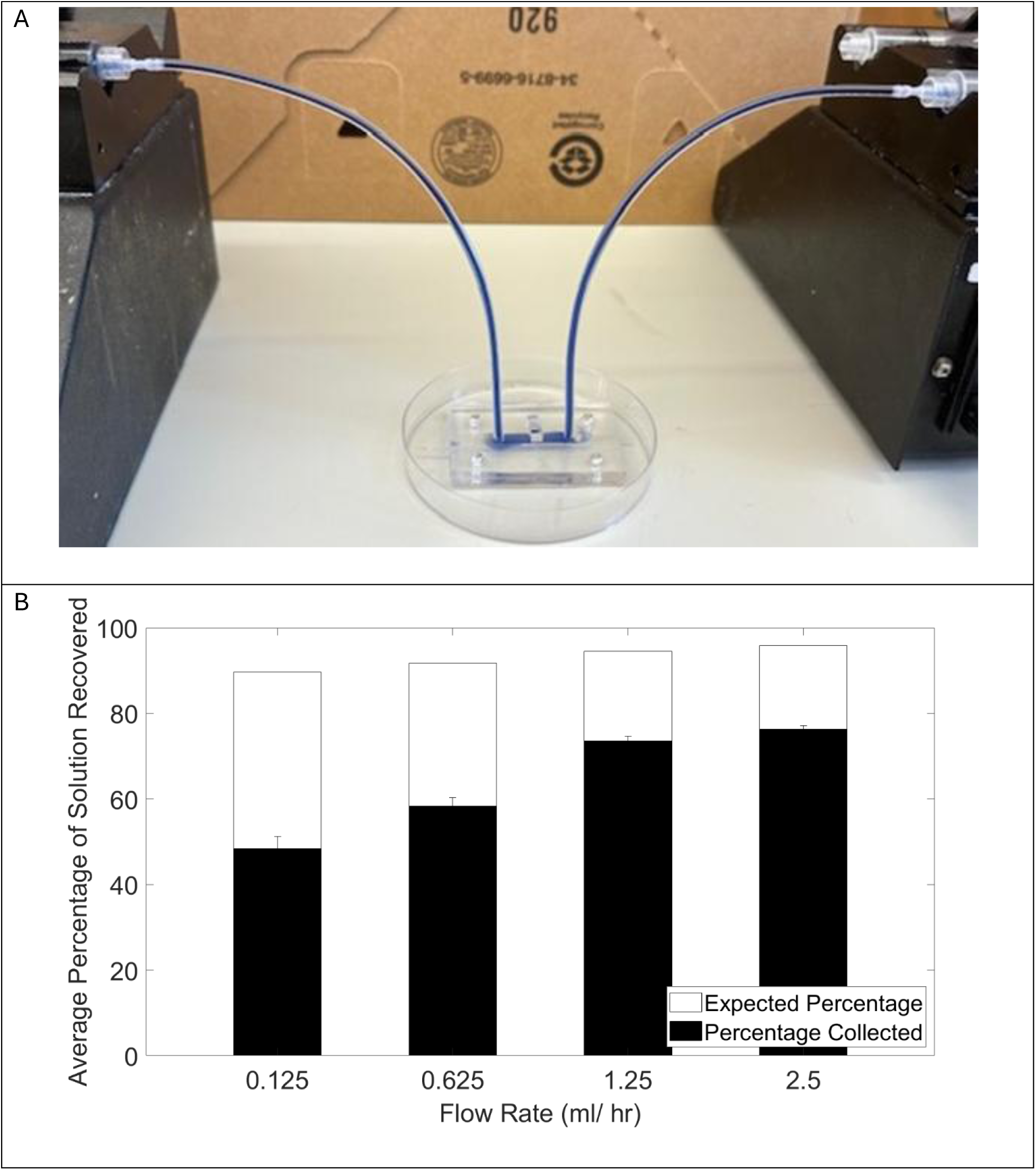
Dyed fluid perfusion testing with early microfluidic platform models. Fluid perfusion testing identified paths media would take during dynamic culture.

**Supplementary Figure 3:**
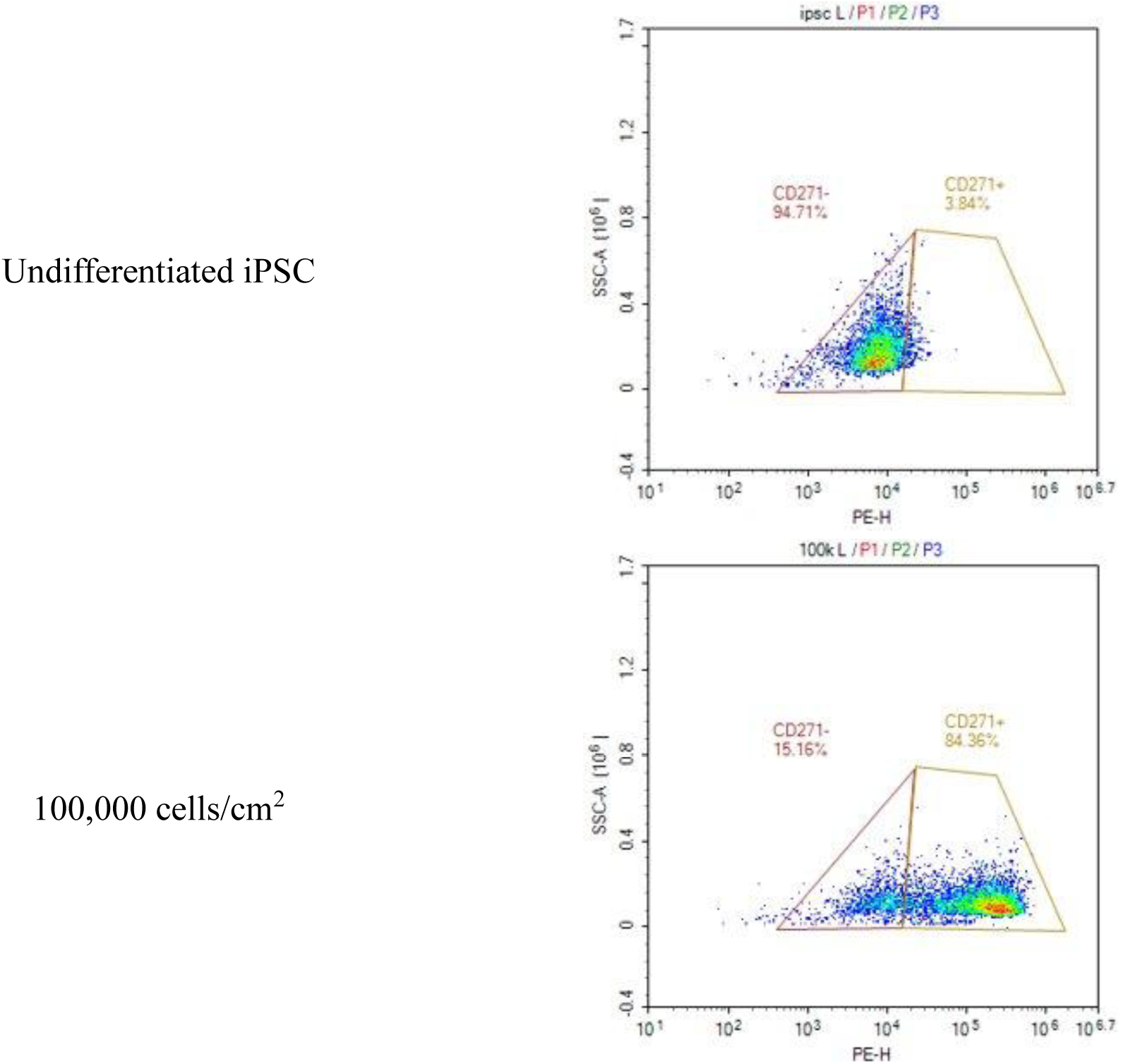
Flow cytometry data for identifying the CD271 neural crest cell (NCC) marker during iPSC differentiation. CD271+ NCCs can be further differentiated into sensory neurons.

**Supplementary Figure 4:**
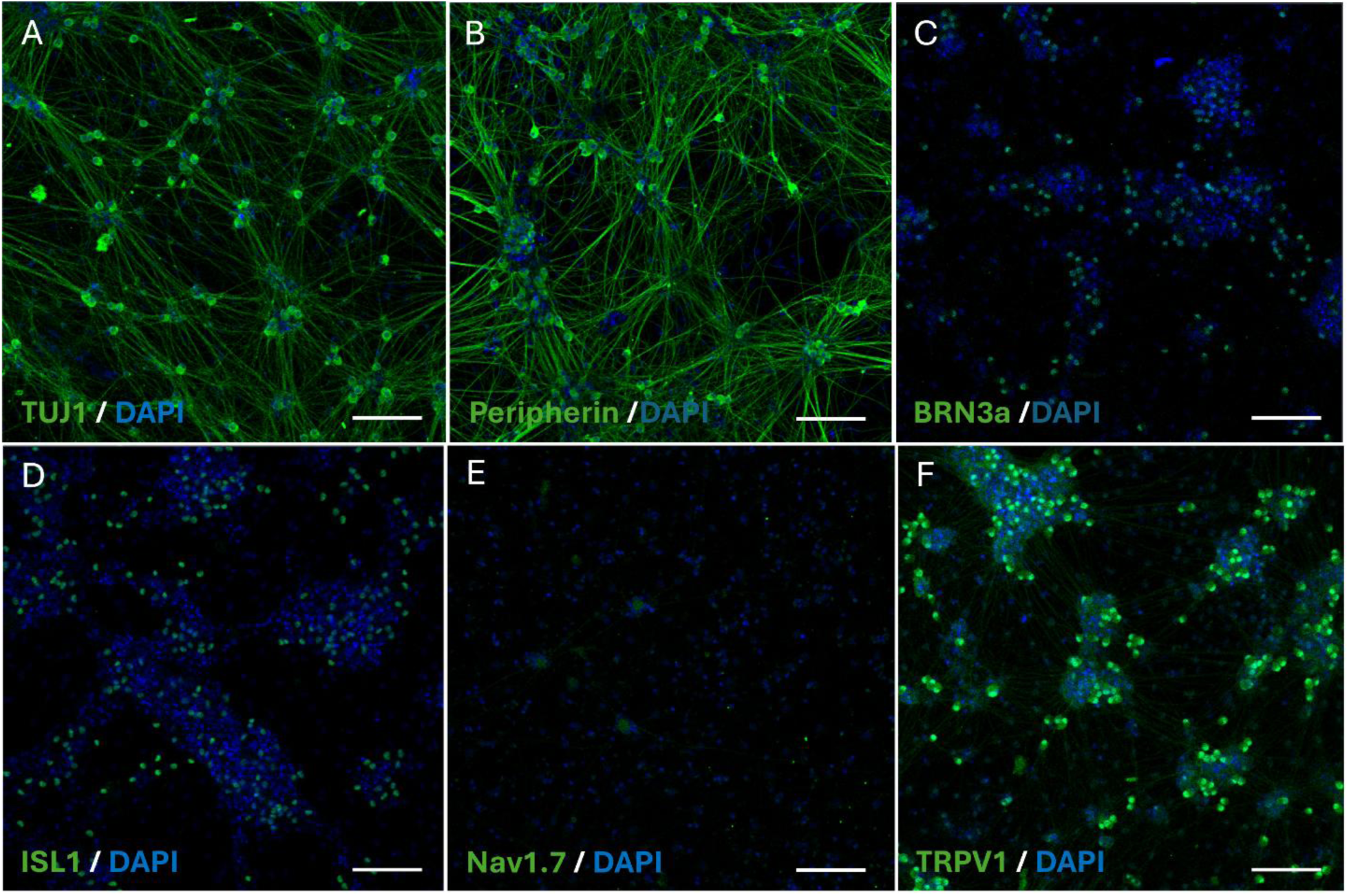
Immunofluorescence analysis confirmed TUJ1, peripherin, BRN3a, ISL1, Na_V_1.7, and TRPV1 expression among SNs differentiated from iPSCs. Images were captured at 20X magnification. Scale bar = 100 μm.

**Supplementary Figure 5:**
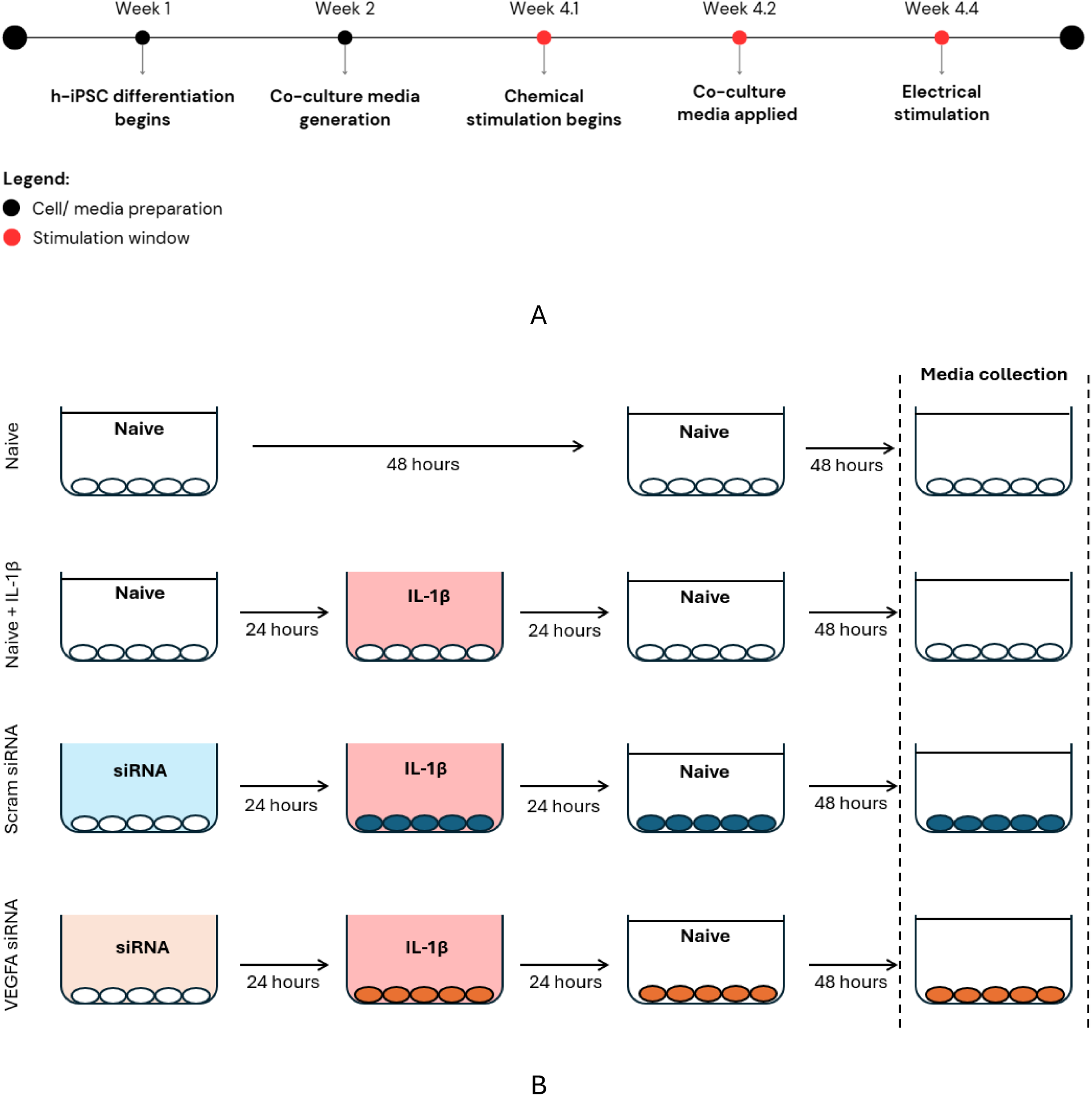
Timelines for donor IVD cell culture and sensory neuron maturation. Approximate 4-week experiment timeline. Mature sensory neurons were cultured with depleted maturation media for three days, two of which included a 1:1 mixture of depleted maturation media and IVD-generated inflammatory media **(A)**. Neurons were then electrically stimulated and Ca^2+^ were assessed. The control group was cultured in naive media (DMEM, 10% FBS, 1% pen-strep) for four days, with a media change after 48 hours. The IL-1β group (not shown) featured IVD cells received an initial 24-hour culture in naive media followed by a 24-hour culture in fresh naive media supplemented by IL-1β (10 ng/mL). A 48-hour culture in fresh naive media concluded this group’s treatment. In the siRNA-treated groups (scram and VEGFA), IVD cells were cultured in alpha MEM media and the appropriate siRNA treatment for the initial 24 hours. This was followed by a 24-hour culture with naive media supplemented with IL-1β. Then, the cells were cultured in fresh naive media for 48 hours **(B)**. Media from each group was collected and frozen at −80 °C after the final 48-hour culture in naive media.

**Supplementary Figure 6:**
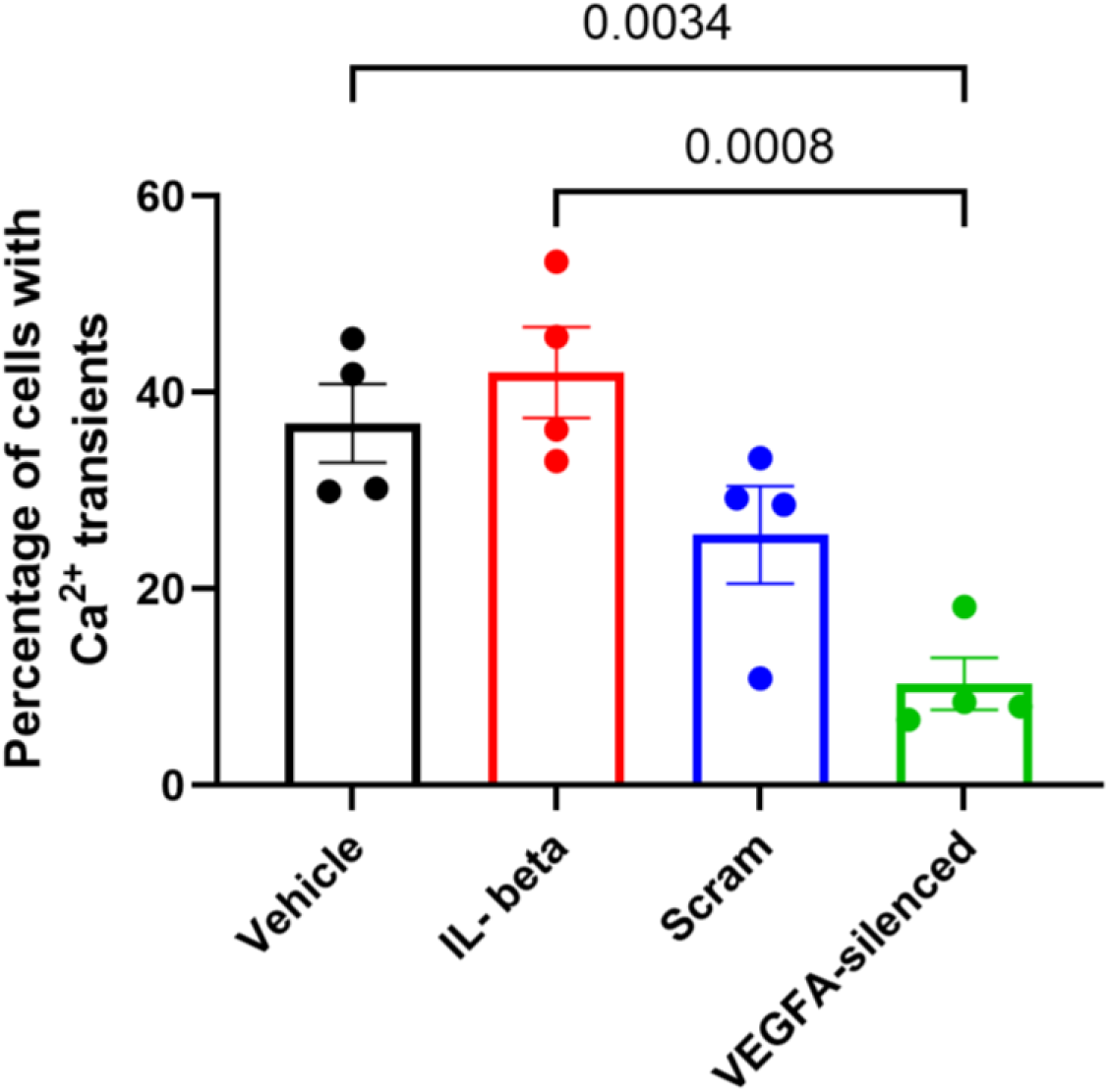
Assessing how SNs exposed to various IVD-generated media responded to EF stimulation. Applying a VEGFA-silencing siRNA pre-treatment to IL-1β-stimulated IVD cells reduced the number of induced Ca^2+^ transient in neurons. Field-wide responses to siRNA treatment revealed muted Ca^2+^ transient levels when compared to the vehicle and IL-1β treatment groups.

**Supplementary Figure 7:**
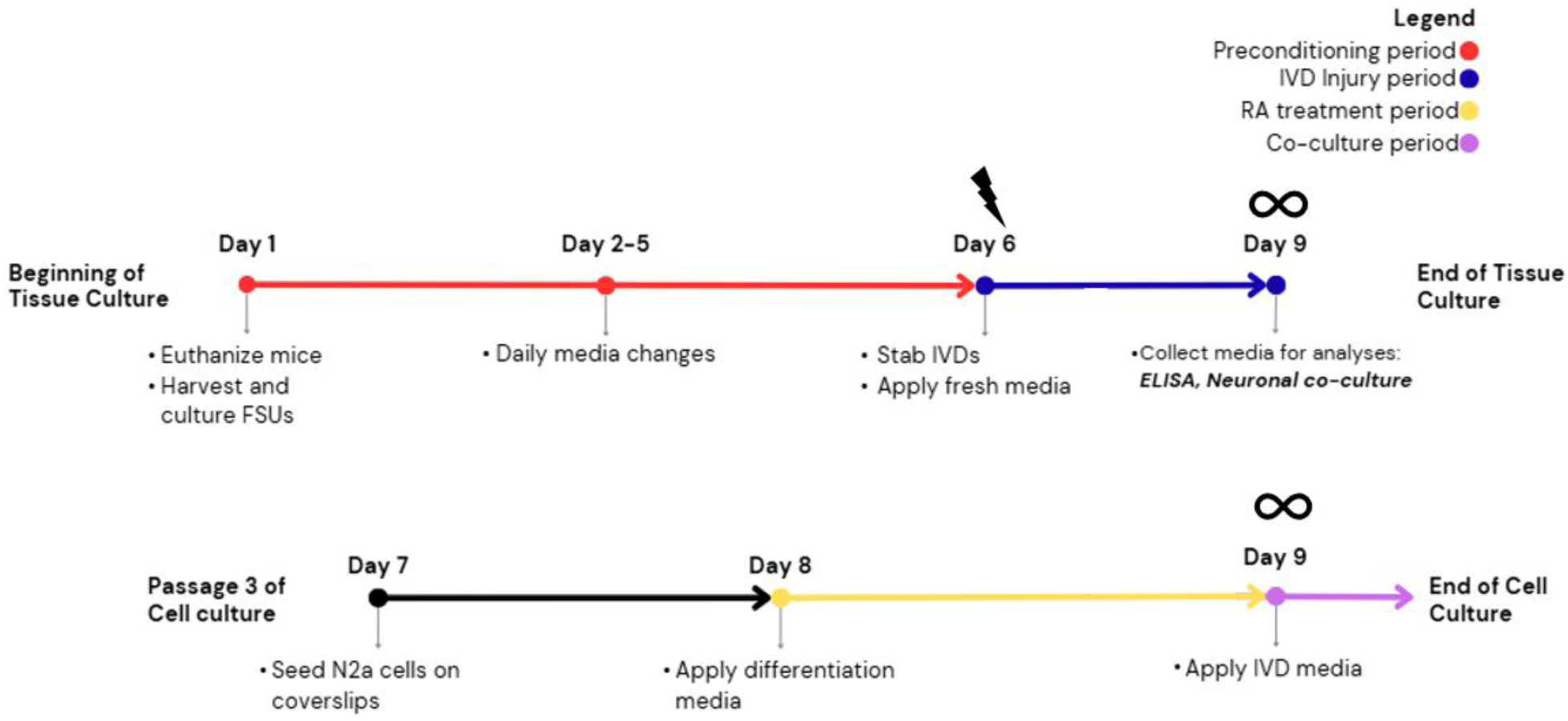
Nine-day culture indirect culture protocol for the murine IVD-SN-like N2a model. Mouse FSUs were preconditioned in tissue culture media (EMEM, 20% FBS, 1% penicillin-streptomycin) for 6 days prior to being injured with a 30G needle and sitting in replenished media for 3 days post-injury without media changes. The 6-day preconditioning period featured daily media changes and inspection of IVD tissue viability. During this period, N2a cells were recovered from cryopreservation and passaged three times before they were cultured on glass coverslips in naive culture media (EMEM, 10% FBS, 1% penicillin-streptomycin) for 24 hours. The cells were then treated with depleted culture media with 10 µM of RA (EMEM, 1% FBS, 1% pen-strep, 10 µM RA) for 24 hours before this media was replaced with IVD-generated media from the tissue culture. After a 24-hour incubation period, the cells were ready for electrophysiology studies.

**Supplementary Figure 8:**
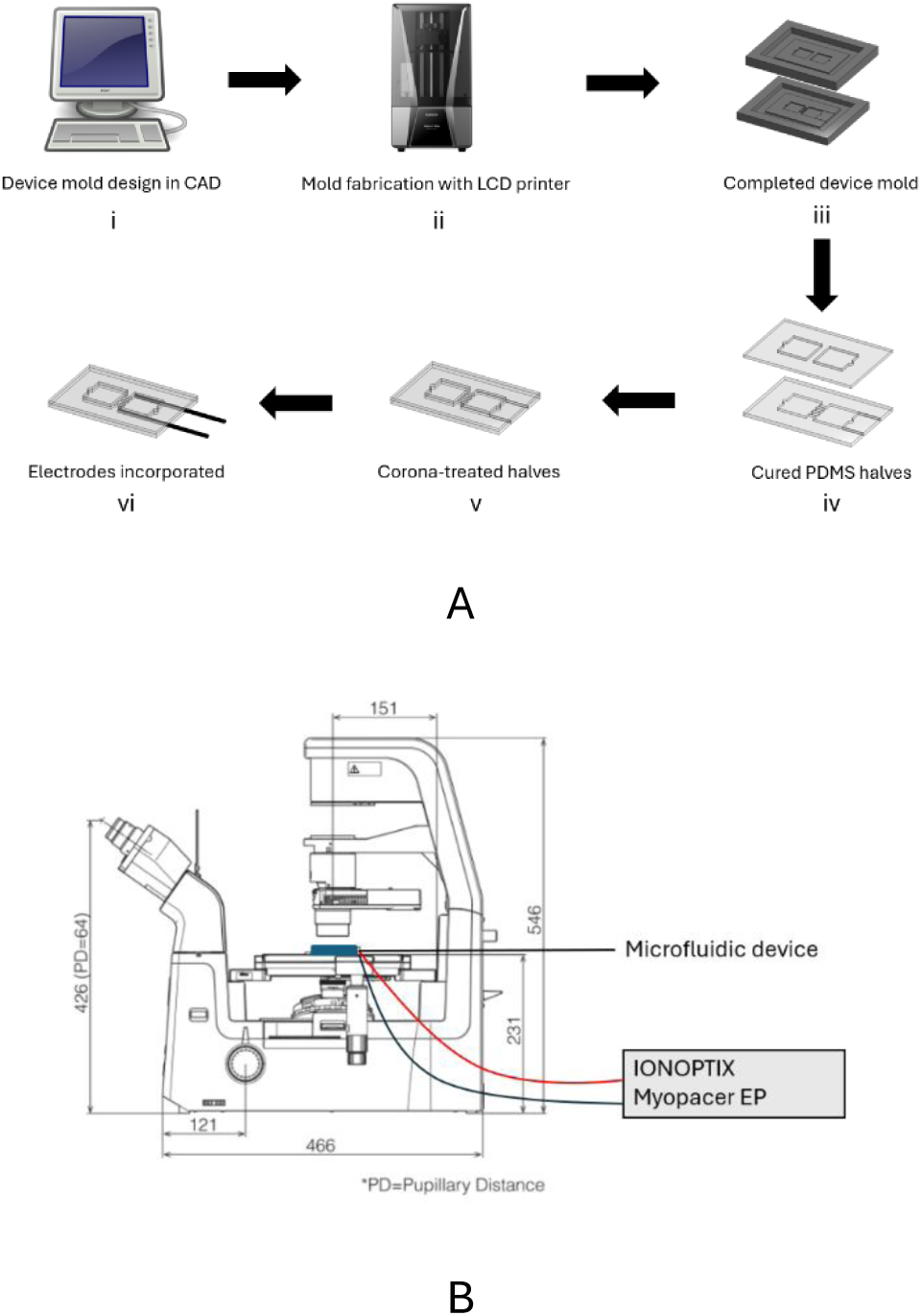
Device fabrication process and incorporation into the experiment apparatus^42^. The resin molds were designed in a 3D computer-aided design (CAD) software **(A.i)** before being 3D printed with an LCD printer **(A.ii)**. A hot 3% agar solution was then poured into the printed molds **(A.iii)** and sent to 4 °C storage for at least 2 hours. Uncured PDMS was poured into the chilled agar molds and cured at 50 °C for 18 hours to form the device’s layers **(A.iv)**. The ‘inner’ faces of both layers were corona-treated, pressed together, and cured for 2 hours at 50 °C **(A.v)**. SS-316L electrodes were then inserted to complete the device build **(A.vi)**. Apparatus of electrical (and chemical) stimulation procedures **(B)**. The microfluidic device (in blue) sits on the microscope platform and wires clipped to its electrodes connect it to a function generator (grey)^15^.

**Supplementary Figure 9:**
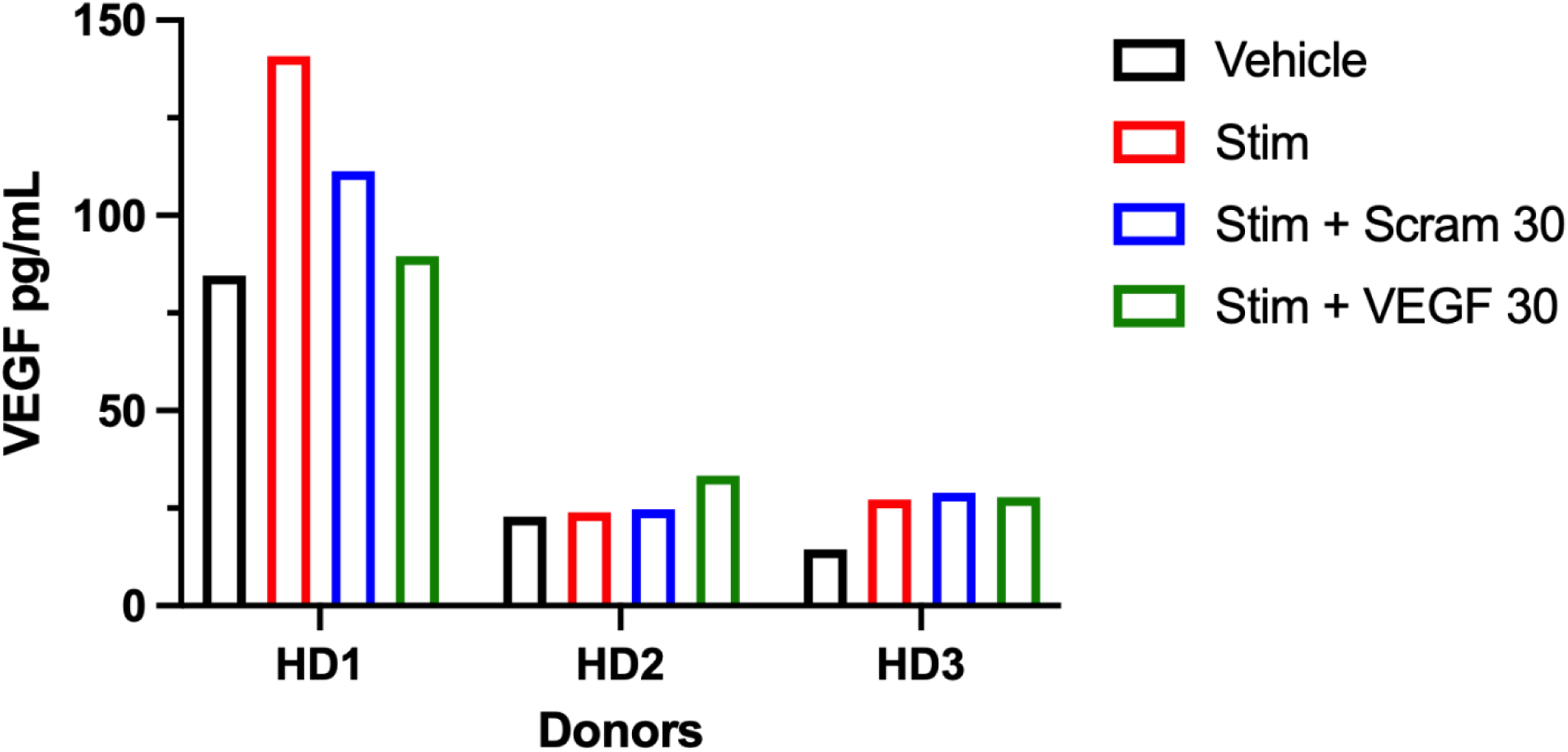
Donor IVD cell responses to IL-1β stimulation after siRNA treatment. Variance in donor response to siRNA VEGF-reduction treatment. Donor 1 (53 y/o female) was more responsive to inflammatory IL-1Β stimulus than donors 2 (73 y/o male) and 3 (70 y/o female).

**Supplementary Table 1.** Material properties assigned to PDMS regions in COMSOL.

| Property | Value | Unit |
| --- | --- | --- |
| Relative permittivity | 2.75 | 1 |
| Density | 970 | kg/m<sup>3</sup> |
| Electric conductivity | $1 \times 10^{-14}$ | S/m |
| Young's modulus | 750 | kPa |
| Poisson's ratio | 0.49 | 1 |

**Supplementary Table 2.** Material properties assigned to water/ culture media regions in COMSOL.

| Property | Value | Unit |
| --- | --- | --- |
| Dynamic viscosity | eta(T) | Pa*s |
| Electrical conductivity | 1.5 | S/m |
| Density | rho(T) | kg/m<sup>3</sup> |
| Relative permittivity | 78 | 1 |

**Supplementary Table 3.** Material properties assigned to SS 430F electrodes in COMSOL.

| Property | Value | Unit |
| --- | --- | --- |
| Electrical conductivity | $1.73913 \times 10^6$ | S/m |
| Relative permittivity | 1 | 1 |
| Density | 7.8 [g/cm<sup>3</sup>] | kg/m<sup>3</sup> |
| Young's modulus | $200 \times 10^9$ | Pa |
| Poisson's ratio | 0.27 | 1 |

**Supplementary Table 4.** Simulation parameters.

| Property | Value |
| --- | --- |
| Inlet flow velocity (m/s) | 0.00106 ms<sup>-1</sup> |
| External applied electric potential (V) | 8 V |

**Supplementary Table 5.** VEGFA production by the human primary IVD cells in response to IL-1Β stimulation after siRNA silencing treatment.

| Age, Gender | Donor | Stim<br>(pg/mL) | No Stim<br>(pg/mL) | Stim + Scram 30<br>(pg/mL) | Stim + VEGFA<br>30<br>(pg/mL) |
| --- | --- | --- | --- | --- | --- |
| 50, F | HD1 | 140.75 | 84.5 | 111.25 | 89.5 |
| 73, M | HD2 | 23.89 | 22.78 | 24.72 | 33.33 |
| 70, F | HD3 | 27.22 | 14.44 | 28.89 | 27.78 |

## References

1. Belitskaya-Levy I, Clark D, Shih M-C, Bair M. Treatment preferences for chronic low back pain: Views of veterans and their providers. Journal of Pain Research. 2021 Jan;Volume 14:161–71. doi:10.2147/jpr.s290400

2. Fatoye F, Gebrye T, Ryan CG, Useh U, Mbada C. Global and regional estimates of clinical and economic burden of low back pain in high-income countries: A systematic review and meta-analysis. Frontiers in Public Health. 2023 Jun 9;11. doi:10.3389/fpubh.2023.1098100

3. U.S. Government Accountability Office. Animal use in research: Federal agencies should assess and report on their efforts to develop and promote alternatives. Washington (DC): GAO; 2019.

4. U.S. Food and Drug Administration. FDA’s roadmap to reducing animal testing. Silver Spring (MD): FDA; 2025 Apr.

5. Lin MTY, et al. Culture of primary neurons from dissociated and cryopreserved mouse trigeminal ganglion. Tissue Eng Part C Methods. 2023;29(8):381–393. doi:10.1089/ten.tec.2023.0054.

6. De Gregorio C, Ezquer F. Sensory neuron cultures derived from adult db/db mice as a simplified model to study type-2 diabetes-associated axonal regeneration defects. Dis Model Mech. 2021;14(1). doi:10.1242/dmm.046334.

7. Potter R, et al. VEGFA-silenced intervertebral disc cells suppress angiogenesis and neurite outgrowth through altered cytokine signaling. J Pain. 2026;41:105773. doi:10.1016/j.jpain.2025.105773.

8. Abraham AC, Liu JW, Tang SY. Longitudinal changes in the structure and inflammatory response of the intervertebral disc due to stab injury in a murine organ culture model. J Orthop Res. 2016;34(8):1431–1438. doi:10.1002/jor.23325.

9. Coriell Institute of Medical Research. GM25256. Available from: https://www.coriell.org/0/Sections/Search/Sample_Detail.aspx?Ref=GM25256&Product=CC (accessed 2026 Mar 2).

10. Clarkson BD, Kahoud RJ, McCarthy CB, Howe CL. Inflammatory cytokine-induced changes in neural network activity measured by waveform analysis of high-content calcium imaging in murine cortical neurons. Sci Rep. 2017;7. doi:10.1038/s41598-017-09182-5.

11. Skaper SD, Facci L, Zusso M, Giusti P. An inflammation-centric view of neurological disease: Beyond the neuron. Front Cell Neurosci. 2018;12. doi:10.3389/fncel.2018.00072.

12. Dai J, et al. Microfluidic disc-on-a-chip device for mouse intervertebral disc: Pitching a next-generation research platform to study disc degeneration. ACS Biomater Sci Eng. 2019;5(4):2041–2051. doi:10.1021/acsbiomaterials.8b01522.

13. Xie W, et al. Intervertebral disc-on-a-chip: A new model for mouse disc culture via integrating mechanical loading and dynamic media flow. Adv Mater Technol. 2023;8(21). doi:10.1002/admt.202300606.

14. Bhatia SN, Ingber DE. Microfluidic organs-on-chips. Nat Biotechnol. 2014;32(8):760–772. doi:10.1038/nbt.2989.

15. Pavesi A, et al. Controlled electromechanical cell stimulation on-a-chip. Sci Rep. 2015;5. doi:10.1038/srep11800.

16. Pavesi A, et al. How to embed three-dimensional flexible electrodes in microfluidic devices for cell culture applications. Lab Chip. 2011;11(9):1593–1595. doi:10.1039/c1lc20084d.

17. Harris J, et al. Fabrication of a microfluidic device for the compartmentalization of neuron soma and axons. J Vis Exp. 2007;(261). doi:10.3791/261.

18. Raj M. PDMS microfluidics: A mini review. J Appl Polym Sci. 2020;137(13). doi:10.1002/app.48958.

19. Toepke MW, Beebe DJ. PDMS absorption of small molecules and consequences in microfluidic applications. Lab Chip. 2006;6(12):1484–1486. doi:10.1039/b612150c.

20. van Meer BJ, et al. Small molecule absorption by PDMS in the context of drug response bioassays. Biochem Biophys Res Commun. 2017;482(2):323–328. doi:10.1016/j.bbrc.2016.11.062.

21. Millet LJ, Stewart ME, Sweedler JV, Nuzzo RG, Gillette MU. Microfluidic devices for culturing primary mammalian neurons at low densities. Lab Chip. 2007;7(8):987–994. doi:10.1039/B705266A.

22. Walk RE, et al. Contrast-enhanced microCT evaluation of degeneration following partial and full width injuries to the mouse lumbar intervertebral disc. Sci Rep. 2022;12(1). doi:10.1038/s41598-022-19487-9.

23. Rohanifar M, et al. Single-cell RNA-sequence analyses reveal uniquely expressed genes and heterogeneous immune cell involvement in the rat model of intervertebral disc degeneration. Appl Sci. 2022;12(16):8244. doi:10.3390/app12168244.

24. Gonzalez CE, et al. The transcriptome and secreted factors of the intervertebral discs in STZ-HFD type 2 diabetic male mice reveal extensive inflammation. bioRxiv. 2024. doi:10.1101/2024.07.31.605332.

25. Clayton SW, et al. Single cell RNA sequencing reveals shifts in cell maturity and function of endogenous and infiltrating cell types in response to acute intervertebral disc injury. bioRxiv. 2024. doi:10.1101/2024.08.10.607363.

26. Walk RE, et al. The neurovascular and inflammatory signatures of the degenerating intervertebral disc. Eur Cells Mater. 2025;54. doi:10.22203/ecm.v054a05.

27. Kumar M, Katyal A. Data on retinoic acid and reduced serum concentration induced differentiation of Neuro-2a neuroblastoma cells. Data Brief. 2018;21:2435–2440. doi:10.1016/j.dib.2018.11.097.

28. Du YJ, et al. Role of miR-124 in the regulation of retinoic acid-induced Neuro-2a cell differentiation. Neural Regen Res. 2020;15(6):1133. doi:10.4103/1673-5374.270417.

29. Schindelin J, et al. Fiji: An open-source platform for biological-image analysis. Nat Methods. 2012;9(7):676–682. doi:10.1038/nmeth.2019.

30. Huebsch N, et al. Automated video-based analysis of contractility and calcium flux in human-induced pluripotent stem cell-derived cardiomyocytes cultured over different spatial scales. Tissue Eng Part C Methods. 2015;21(5):467–479. doi:10.1089/ten.tec.2014.0283.

31. Warner Instruments. Perfusion chamber with field stimulation (RC-49MFSH). Available from: https://www.warneronline.com/perfusion-chamber-with-field-stimulation-rc-49mfsh (accessed 2025 Dec 24).

32. Biagini F, et al. A millifluidic chamber for controlled shear stress testing: Application to microbial cultures. Ann Biomed Eng. 2023;51(12):2923–2933. doi:10.1007/s10439-023-03361-4.

33. Visone R, et al. A microscale biomimetic platform for generation and electro-mechanical stimulation of 3D cardiac microtissues. APL Bioeng. 2018;2(4). doi:10.1063/1.5037968.

34. Mainardi A, et al. Intervertebral disc-on-a-chip as advanced in vitro model for mechanobiology research and drug testing: A review and perspective. Front Bioeng Biotechnol. 2022;9. doi:10.3389/fbioe.2021.826867.

35. Weng X, et al. Chronic inflammatory pain is associated with increased excitability and hyperpolarization-activated current (IH) in C- but not Aδ-nociceptors. Pain. 2012;153(4):900–914. doi:10.1016/j.pain.2012.01.019.

36. Odem MA, Bavencoffe AG, Cassidy RM, Lopez ER, Tian J, Dessauer CW, et al. Isolated nociceptors reveal multiple specializations for generating irregular ongoing activity associated with ongoing pain. Pain. 2018 Jul 12;159(11):2347–62. doi:10.1097/j.pain.0000000000001341

37. Bavencoffe A, Lopez ER, Johnson KN, Tian J, Gorgun FM, Shen BQ, et al. Widespread hyperexcitability of Nociceptor somata outlasts enhanced avoidance behavior after incision injury. Pain. 2024 Oct 22;166(5):1088–104. doi:10.1097/j.pain.0000000000003443

38. Xie W, Xing Y, Xiao L, Zhang P, Oh R, Zhang Y, et al. Intervertebral disc-on-a-chipmf: A new model for mouse disc culture via integrating mechanical loading and dynamic media flow. Advanced Materials Technologies. 2023 Aug 27;8(21). doi:10.1002/admt.202300606

39. Pattappa G, Peroglio M, Sakai D, Mochida J, Benneker L, Alini M, et al. CCL5/Rantes is a key chemoattractant released by degenerative intervertebral discs in organ culture. European Cells and Materials. 2014 Feb 6;27:124–36. doi:10.22203/ecm.v027a10

40. Xiong C, et al. Human induced pluripotent stem cell derived sensory neurons are sensitive to the neurotoxic effects of paclitaxel. Clin Transl Sci. 2020;14(2):568–581. doi:10.1111/cts.12912.

41. Gold MS, Flake NM. Inflammation-mediated hyperexcitability of sensory neurons. Neurosignals. 2005;14(4):147–157. doi:10.1159/000087653.

42. Torsney C. Inflammatory pain neural plasticity. Curr Opin Physiol. 2019;11:51–57. doi:10.1016/j.cophys.2019.06.001.

43. Zhang YH, et al. Editorial: Inflammatory pain: Mechanisms, assessment, and intervention. Front Mol Neurosci. 2023;16. doi:10.3389/fnmol.2023.1286215.

44. Nikon Instruments Inc. ECLIPSE Ts2R inverted microscope specifications. 2024. Available from: https://www.nikon.com

45. Chan CY, et al. Accelerating drug discovery via organs-on-chips. Lab Chip. 2013;13(24):4697. doi:10.1039/c3lc90115g.

46. Bhise NS, et al. Organ-on-a-chip platforms for studying drug delivery systems. J Control Release. 2014;190:82–93. doi:10.1016/j.jconrel.2014.05.004.

47. Berke IM, et al. Electric field stimulation for the functional assessment of isolated dorsal root ganglion neuron excitability. Ann Biomed Eng. 2021;49(3):1110–1118. doi:10.1007/s10439-021-02725-y.

48. Zaszczyńska A, et al. Piezoelectric scaffolds as smart materials for bone tissue engineering. Polymers. 2024;16(19):2797. doi:10.3390/polym16192797.

49. Yue W, et al. The role of piezoelectric materials in bone remodeling and repair: Mechanisms and applications. Int J Nanomed. 2025;20:11593–11616. doi:10.2147/ijn.s535976.

50. Neužil P, et al. Revisiting lab-on-a-chip technology for drug discovery. Nat Rev Drug Discov. 2012;11(8):620–632. doi:10.1038/nrd3799.

51. Meng X, Yu Y, Jin G. Numerical simulation and experimental verification of droplet generation in microfluidic digital PCR chip. Micromachines. 2021;12(4):409. doi:10.3390/mi12040409.

52. Hernández-Cid D, et al. Modeling droplet formation in microfluidic flow-focusing devices using the two-phase level set method. Mater Today Proc. 2022;48:30–40. doi:10.1016/j.matpr.2020.09.417.

53. Jebari N, et al. 3D simulation-driven design of a microfluidic immunosensor for real-time monitoring of sweat biomarkers. Micromachines. 2024;15(8):936. doi:10.3390/mi15080936.

54. Merck KGaA. MilliporeSigma life science products. 2024. Available from: https://www.emdmillipore.com/US/en

55. Ibidi GmbH. μ-Slide I Luer: Channel slide for flow assays. 2024. Available from: https://ibidi.com/channel-slides/50--slide-i-luer.html

56. Flexcell International Corp. FlexFlow shear stress microscopy. Available from: https://www.flexcellint.com/products/flexflow-r-microscopy (accessed 2025 Dec 24).

57. Comsol AB. COMSOL Multiphysics version 6.3. Stockholm: COMSOL; 2024. Available from: https://www.comsol.com

58. Plonsey R, Barr RC. Bioelectricity: A Quantitative Approach. 3rd ed. New York: Springer; 2007.

59. Keener J, Sneyd J. Mathematical Physiology. 2nd ed. New York: Springer; 2009.

60. Microsoft Corporation. Microsoft Excel version 365. Redmond (WA); 2024.

61. GraphPad Software. GraphPad Prism version 11.0.0. San Diego (CA); 2024.

62. Lindsay RM. Role of neurotrophins and Trk receptors in the development and maintenance of sensory neurons: An overview. Philos Trans R Soc Lond B Biol Sci. 1996;351(1338):365–373. doi:10.1098/rstb.1996.0030.

63. Kirstein M, Fariñas I. Sensing life: Regulation of sensory neuron survival by neurotrophins. Cell Mol Life Sci. 2002;59(11):1787–1802. doi:10.1007/pl00012506.

64. Jones I, et al. Development and validation of an in vitro model system to study peripheral sensory neuron development and injury. Sci Rep. 2018;8. doi:10.1038/s41598-018-34280-3.

65. Lin MTY, et al. Culture of primary neurons from dissociated and cryopreserved mouse trigeminal ganglion. Tissue Eng Part C Methods. 2023;29(8):381–393. doi:10.1089/ten.tec.2023.0054.

66. Sochocka M, Diniz BS, Leszek J. Inflammatory response in the CNS: Friend or foe? Mol Neurobiol. 2016;54(10):8071–8089. doi:10.1007/s12035-016-0297-1.

67. Counil H, Krantic S. Synaptic activity and (neuro)inflammation in Alzheimer’s disease: Could exosomes be an additional link? J Alzheimers Dis. 2020;74(4):1029–1043. doi:10.3233/jad-191237.

68. Teixeira I, et al. Polydimethylsiloxane mechanical properties: A systematic review. AIMS Mater Sci. 2021;8(6):952–973. doi:10.3934/matersci.2021058.

